# Deep deconvolution of the hematopoietic stem cell regulatory microenvironment reveals a high degree of specialization and conservation between mouse and human

**DOI:** 10.1101/2021.07.17.452614

**Authors:** Jin Ye, Isabel A. Calvo, Itziar Cenzano, Amaia Vilas, Xabier Martinez-de-Morentin, Miren Lasaga, Diego Alignani, Bruno Paiva, Ana C. Viñado, Patxi San Martin-Uriz, Juan P. Romero, Delia Quilez Agreda, Marta Miñana Barrios, Ignacio Sancho González, Gabriele Todisco, Luca Malcovati, Nuria Planell, Borja Saez, Jesper Tegner, Felipe Prosper, David Gomez-Cabrero

## Abstract

Understanding the regulation of normal and malignant human hematopoiesis requires comprehensive cell atlas of the hematopoietic stem cell (HSC) regulatory microenvironment. Here, we develop a tailored bioinformatic pipeline to integrate public and proprietary single-cell RNA sequencing (scRNA-seq) datasets. As a result, we robustly identify for the first time 14 intermediate cell states and 11 stages of differentiation in the endothelial and mesenchymal BM compartments, respectively. Our data provide the most comprehensive description to date of the murine HSC-regulatory microenvironment and suggests a higher level of specialization of the cellular circuits than previously anticipated. Furthermore, this deep characterization allows to infer conserved features in human, suggesting that the layers of microenvironmental regulation of hematopoiesis may also be shared between species. Our resource and methodology are a steppingstone towards a comprehensive cell atlas of the BM microenvironment.

## INTRODUCTION

Uncovering pathogenetic mechanisms requires identifying the corresponding major groups of genes in the disease-relevant tissues(Gomez-Cabrero et al., 2014; Wolkenhauer et al., 2013). To this end, collective efforts such as the Human Single-Cell Atlas have been launched, aiming at providing a single-cell map of human tissues and organs. A core case is the hematopoietic system, where single-cell RNA sequencing (scRNA-seq) has allowed to refine our understanding of hematopoiesis in mouse and human(Giladi et al., 2018; Nestorowa et al., 2016; Rodriguez-Fraticelli et al., 2018). Furthermore, these studies have challenged the classical view of hematopoiesis differentiation as a compendium of discrete cellular states with decreased differentiation potential towards a more dynamic view in which hematopoietic stem and progenitor cells (HSPC) gradually pass through a continuum of differentiation states(Karamitros et al., 2017; Laurenti and Göttgens, 2018; Velten et al., 2017; Weinreb et al., 2020). Moreover, recent studies using scRNA-seq technologies have shed light on the organization of the hematopoietic regulatory microenvironment in the mouse(Baccin et al., 2020; Baryawno et al., 2019; Kanazawa et al., 2021; Matsushita et al., 2020; Tikhonova et al., 2019; Wolock et al., 2019; Zhong et al., 2020). These studies have resolved some of the controversies regarding the overlap of stromal populations previously described and the description of certain discrete stromal cells as professional, hematopoietic cytokine-producing populations(Baccin et al., 2020). Moreover, further combination with *in-situ* technologies helped to delineate the relationship between specific stromal cell types in the murine Bone marrow (BM)(Baccin et al., 2020). This data has provided a wider and more dynamic picture of hematopoiesis and their regulatory microenvironment, allowing for a provocative hypothesis to rise, such as whether their specific association with given niches controls transcriptional states in hematopoietic stem cells and whether these states are reversible upon occupying alternative niches.

Nevertheless, these studies are limited by the number of cells sequenced, potentially hampering our ability to resolve the full spectrum of cellular states and differentiation stages that define the stromal BM microenvironment. Further, knowledge on the conservation of the cellular composition in the human BM stroma is in its infancy due to the difficulty of obtaining high-quality samples with sufficient stromal cell numbers from healthy individuals. This leaves us with two outstanding challenges; how to piece together such different fragments towards a comprehensive molecular atlas and to what extent such an atlas in mice is conserved in the human bone marrow.

Here, we integrate three scRNA-seq datasets (two publicly available(Baryawno et al., 2019; Tikhonova et al., 2019) and one in-house) separately targeting two well-defined populations (endothelial and mesenchymal cells). The integration of distinct data sets required developing tailored bioinformatics pipelines to ensure the robust identification of cell types and stages. We identify 14 endothelial subclusters and 11 subpopulations defining different stages of differentiation in the mesenchyme. Our analysis provides the most comprehensive atlas of the cellular composition in the mouse bone marrow. Last, we asked to what extent such an atlas could provide insight into the less accessible human BM microenvironment. To this end, we made the first pilot study, profiling the human BM using scRNA-seq, which was integrated with our mouse BM atlas. This analysis demonstrated substantial conservation between species.

## RESULTS

### Data integration and high-resolution clustering strategy

Figure 1 provides a graphical summary of the experimental design and the analysis flow. We integrated selected subsets of cells from three distinct mouse datasets: two recently published (“Tikhonova et al.”(Tikhonova et al., 2019), 6626 cells, and “Baryawno et al.”(Baryawno et al., 2019), 38443 cells) and an independent dataset (“*In-house*” dataset, 13402). These datasets differ in the procedures for isolation of cells within the BM microenvironment. This includes unbiased isolation of cells lacking hematopoietic markers (“*Baryawno*”(Baryawno et al., 2019) and “*In-house*”) **(Fig. S1a)** versus targeted isolation of populations of interest as in Tikhonova et al.(Tikhonova et al., 2019). Furthermore, not every cell type identified in one study is present in the other datasets.

**Figure 1.**
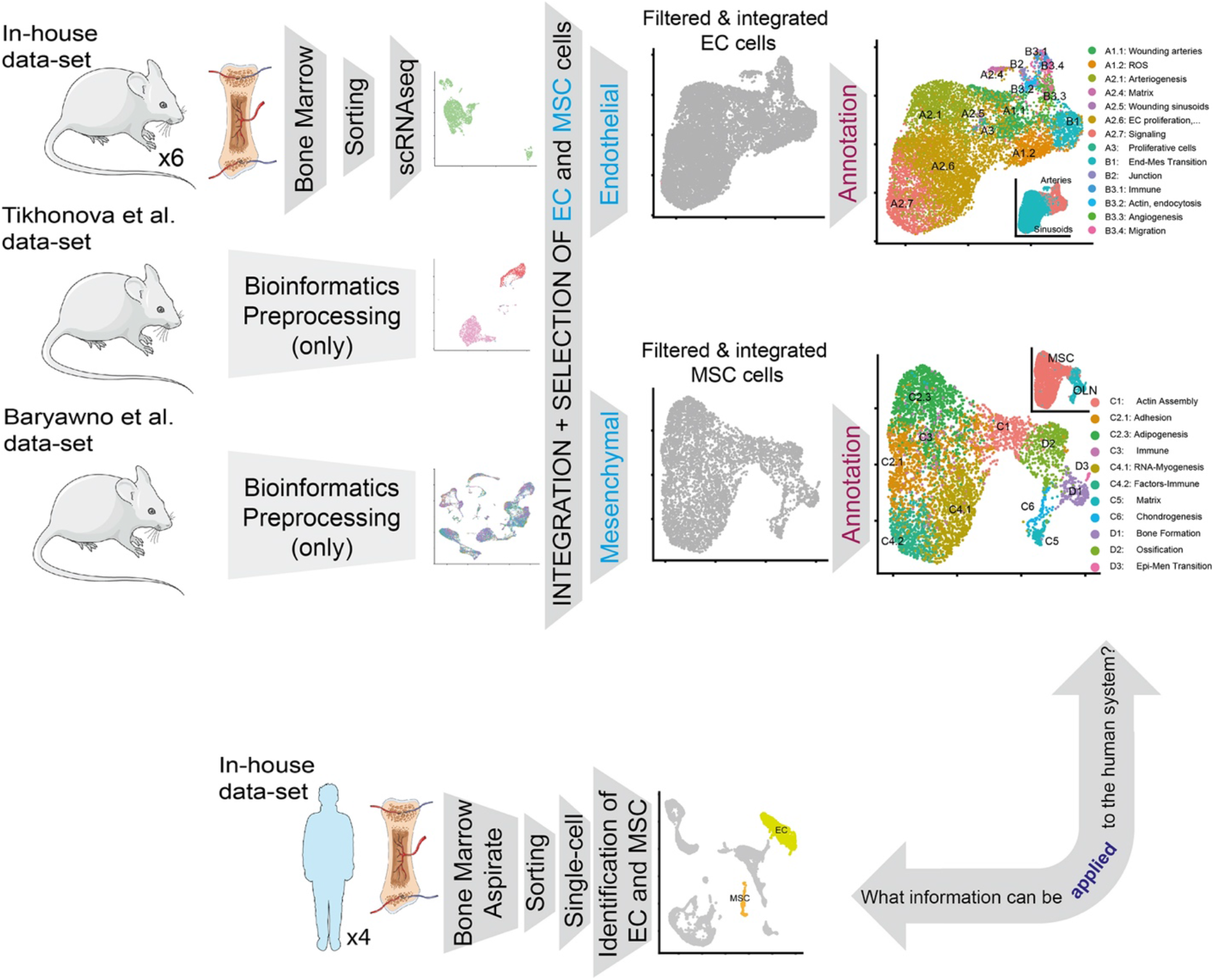
Overview of the paper. Graphical brief description of the paper.

We decided to focus our analysis on those bona fide niche populations such as endothelial (EC) and mesenchymal (MSC) cells due to their presence in the three studies and their relevance in controlling **hematopoietic stem cell** (HSC) maintenance. For the integrative analysis, we used the Tikhonova study as a reference to facilitate the integration, considering that their cells were isolated based on fluorescence reporter expression driven by cell-specific gene promoters: VE-Cad for endothelial cells and LEPR for mesenchymal cells. Therefore, to identify and label the cells of interest, we integrated separately “In house + Tikhonova” and “Baryawno + Tikhonova” **(Fig. 2a)** using the cell labels from the Tikhonova study. As a result, and after quality filters (see Methods), we labeled in each dataset endothelial cells (N=9587) and mesenchymal cells (N=5291) that were used for integration of the three datasets for each cell-type.

**Figure 2.**
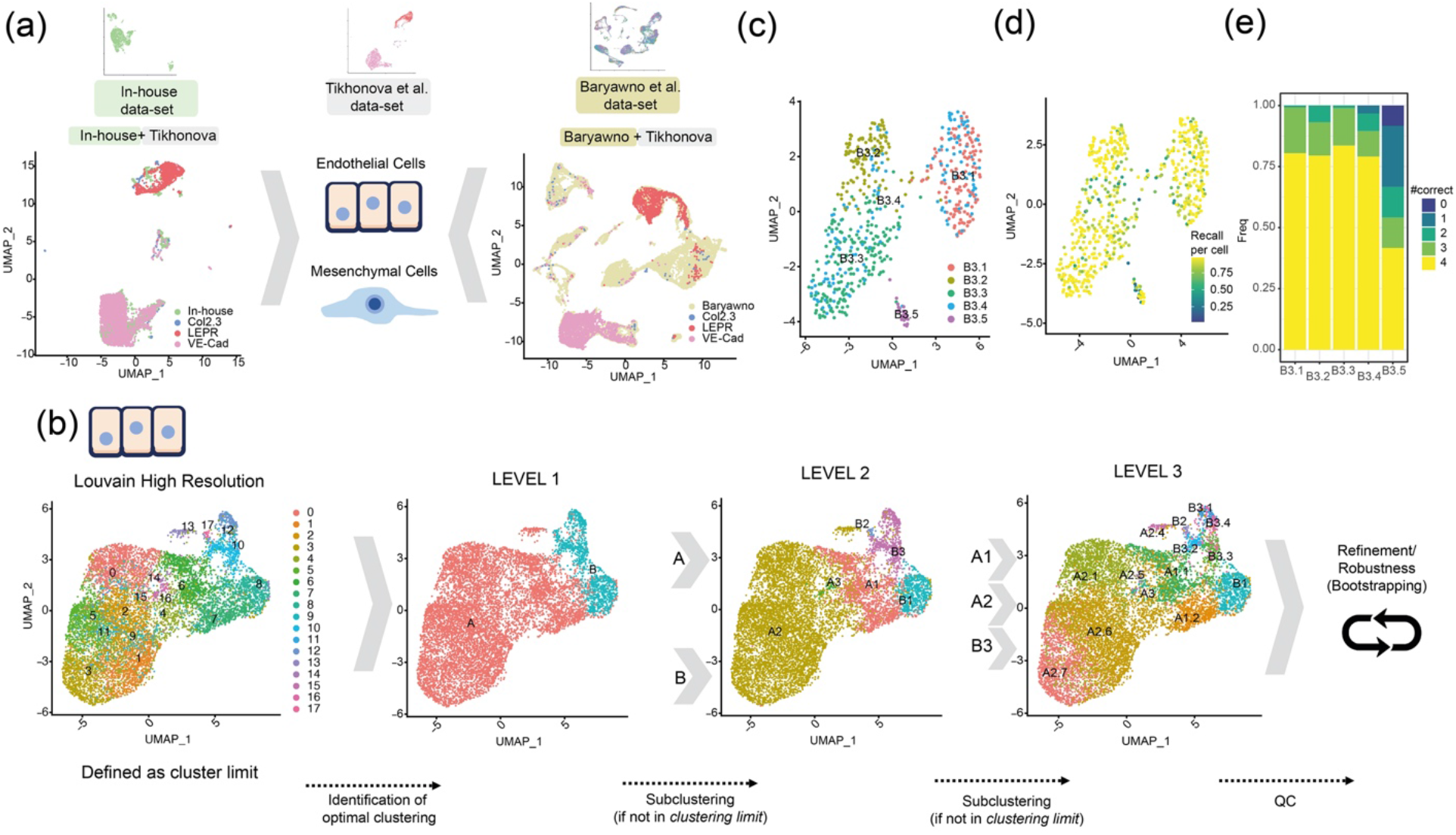
Data integration and high-resolution clustering strategy. **(a)** Integrated analysis of the bone marrow niche datasets (two publicly available, Tikhonova et al. and Baryawno et al., and one generated in our lab, *in-house*) separately for two well-defined populations (endothelial and mesenchymal cells). Tikhonova et al. data set are used as a reference considering their separated cell profiling strategy for COL2.3+, LEPR+ and VE-Cad+ populations. In the top row, a UMAP projection is depicted for each single-cell RNA dataset. In the left-lower (“In-house + Tikhonova”) and in the right-lower (“Baryawno + Tikhonova”) data sets are integrated to identify endothelial and mesenchymal populations. **(b)** Clustering strategy: the analysis of endothelial cells as an example. An upper limit to the cluster is set for the clustering (left panel) using Louvain high-resolution clustering. Then, an iterative divide-and-conquer strategy identifies the optimal level of clusters at different levels: Level 1 (second panel from the left), Level 2 (third panel), and Level 3 (fourth panel). **(c), (d), (e)** The robustness analysis for sub-clustering B3 (from Level 2 to Level 3). Specifically: **(c)** sub-clusters identified, **(d)** the fraction of assignments to its original cluster *using a random-forest + bootstrapping strategy* and **(e)** summary of the results (d) per cluster, #correct indicates the times a cluster is a dominant cluster (see Methods) for a cell within in all pairwise comparisons (see Fig.S2 for the sub-clustering analysis of A1 and A2).

Following the integration, endothelial and mesenchymal groups were used to identify cellular subtypes and stage-specific cells **(Fig. 2, Fig. S2 and S3)**. However, current state-of-the-art clustering methodologies, including Louvain clustering(Blondel et al., 2008; Traag et al., 2019), could not discriminate robustly among different cell subtypes **(Fig. 2b left panel and Fig. S3a)**, in part because there is a high degree of cell-to-cell similarity when considering cells of the same origin(Tasic et al., 2016a). To enable robust sub-clustering, we customized an existing bootstrapping-based approach. In brief, first, a divide-and-conquer strategy is applied, where the first level of robust clusters is identified (see Methods). Next, we proceeded with another round of clustering, yielding the second level of robust sub-clusters. As a termination criterion, no sub-clustering was considered if a cluster was found to have no sub-divisions in the Louvain high-resolution clustering(Blondel et al., 2008; Macosko et al., 2015) **(Fig. 2b left panel and Fig. S3a)**. As a result, the cells are grouped into clusters; then, in the second step, we applied a bootstrapping-based methodology adapted from the Bosiljka study(Tasic et al., 2016b) (see Methods and **Fig. S2a)** to quantify the robustness of each cluster. We formulated two evaluation metrics: for each cell, we computed “*how many times it has been correctly assigned to the cluster proposed*” (e.g. recall per cell in Fig.S2d), and for each cluster, we quantified “how many times a cluster remains dominant (#Correct) in all comparisons for cells within it” (e.g. #correct in Fig.S2e); see Methods for a detailed explanation. If a non-robust cluster was identified, the cells of such cluster were then assigned to the neighboring clusters repeating the Random-Forest based strategy. For instance, in the analysis of ECs, three levels of sub clustering were conducted (**Fig. 2b** middle and right panels). In Level 1, two clusters were identified (A and B), and considering that possible sub-clusters were identified in the Louvain high-resolution clustering **(Fig.2b**, (second panel from the left), they were sub-clustered resulting in Level 2. After Level 2 only three subclusters were further investigated (A1, A2, B3) in Level 3 **(Fig.2b, third panel)**. Level 3 was identified as the final level, and a robustness analysis was conducted for the Level 3 clusters **(Fig. 2b fourth panel, c-e and Fig. S2c-h)**. Non-robust clusters were eliminated and their cells were reassigned to neighboring clusters (see Methods). Similarly, we applied the same robust clustering to the mesenchymal stromal cells **(Fig. S3a-h)**. By using this approach, we were able to describe 14 subclusters in the endothelium **(Fig. 2b fourth panel and Fig. 3a)** and 11 in the mesenchyme **(Fig. S3a fourth panel and Fig. 3c)**. Encouragingly, we observed that the final subclusters were not biased to any study or any specific cell stage **(Fig. 3b,d and Fig. S2b)** and only clusters B2 and D3 did not have cells from the three datasets.

**Figure 3.**
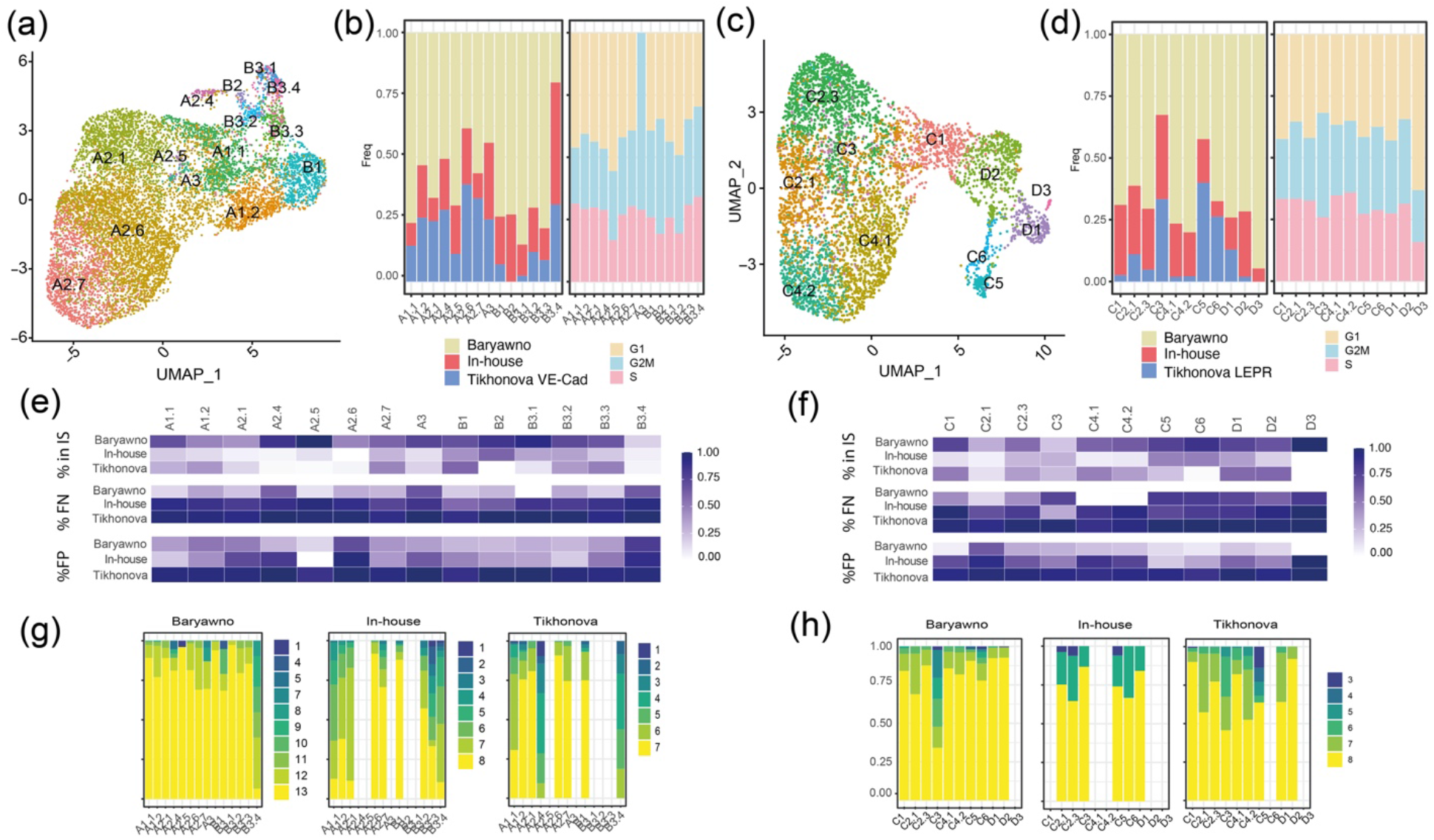
Quality control and added value of the clustering analysis. **(a)** Representation of the final endothelial clusters. **(b)** Left panel: proportion of cells per dataset in each cluster in the final Endothelial clustering analysis. Right panel: proportion of cells per cell cycle stage using Seurat in each cluster in the final Endothelial clustering analysis. **(c**,**d)** Similar as (a,b) for MSC. **(e**,**f)** The added value of the integrated approach. Upper panel: every cell depicts the % of markers identified per cluster using only one dataset when compared with the markers identified in the total dataset. Middle panel: every cell depicts the % of False Negatives. Bottom panel: every cell depicts the % of False Positives when comparing the analysis conducted within each dataset with the integrated analysis (considered as the correct result). (e) and (f) are respectively associated with endothelial and mesenchymal cells. **(g**,**h)** Robustness of the cluster characterization using only cells from a single dataset but maintaining the same cluster structure. (g) and (h) are respectively associated with endothelial and mesenchymal cells.

By integrating the three datasets, distinct cellular states in the microenvironment and the description of the gene markers defining those subtypes were identified. The rationale being that despite the partial overlap between the datasets, a larger number of cells would generally contribute to a deeper characterization of the subpopulations. To directly address this and using the markers derived from the integrative analysis, we conducted a series of analyses to assess whether the integration provided additional insights compared to each dataset.

First, we investigated what percentage of final integrated-based markers were recovered by each dataset by itself **(Fig. 3e,f upper panel)**, what percentage of false negatives **(Fig. 3e,f middle panel)**, and false positives **(Fig. 3e,f lower panel)**. For some of the subclusters, such as B3.4, A2.1, and A2.6 (endothelium) or C2.1 and C3 (mesenchyme), over 50% of the markers could not be detected by each dataset separately. In a second analysis, we quantified the robustness of the defined clusters using data from each study separately **(Fig. 3g,h)**. Only the Baryawno dataset allows for the robust identification of all the subclusters except for B3.4 in the endothelium. However, performing the same clustering strategy using only the Baryawno’s dataset in the endothelium, could not identify all the subclusters with the same level of resolution compared to those observed with the integrated dataset **(Fig. S4a)**.

Taken together, these data demonstrate that our customized approach for the integration of multiple datasets allows for a robust deconvolution of cell states when there is a high degree of similarity between cells of the same origin.

### Deep characterization of the BM endothelial cell compartment

Next, we aimed to characterize each of those 14 endothelial subclusters **(Fig. 3a)** based on the identified markers **(Table S1)**. Using the expression of those molecular markers, we could discriminate between arteries and sinusoids **(Fig. 4a and Table S2)** in agreement with previous reports(Hooper et al., 2009; Itkin et al., 2016; Rafii et al., 2016). Arterial clusters showed high expression of specific arterial genes such as *Ly6a, Ly6c1, Igfbp3*, and *Vim* **(Fig. 4b upper panels)**. At the same time, sinusoidal cells were defined by their characteristic signature expressing *Adamts5, Stab2, Il6st, and Ubd* **(Fig. 4b lower panels)**. Importantly, besides these already known markers, differential expression analysis of the integrative datasets revealed some novel genes specific of each endothelial population, such as *Igfbp7* and *Ppia* for arteries and *Cd164* or *Blrv* for sinusoids **(Fig. 4c)**. The expression of these genes would be consistent with the role *Igfbp7* and *Blvrb* in the maintenance of endothelial vasculature homeostasis(Klóska et al., 2019; Tamura et al., 2009).

**Figure 4.**
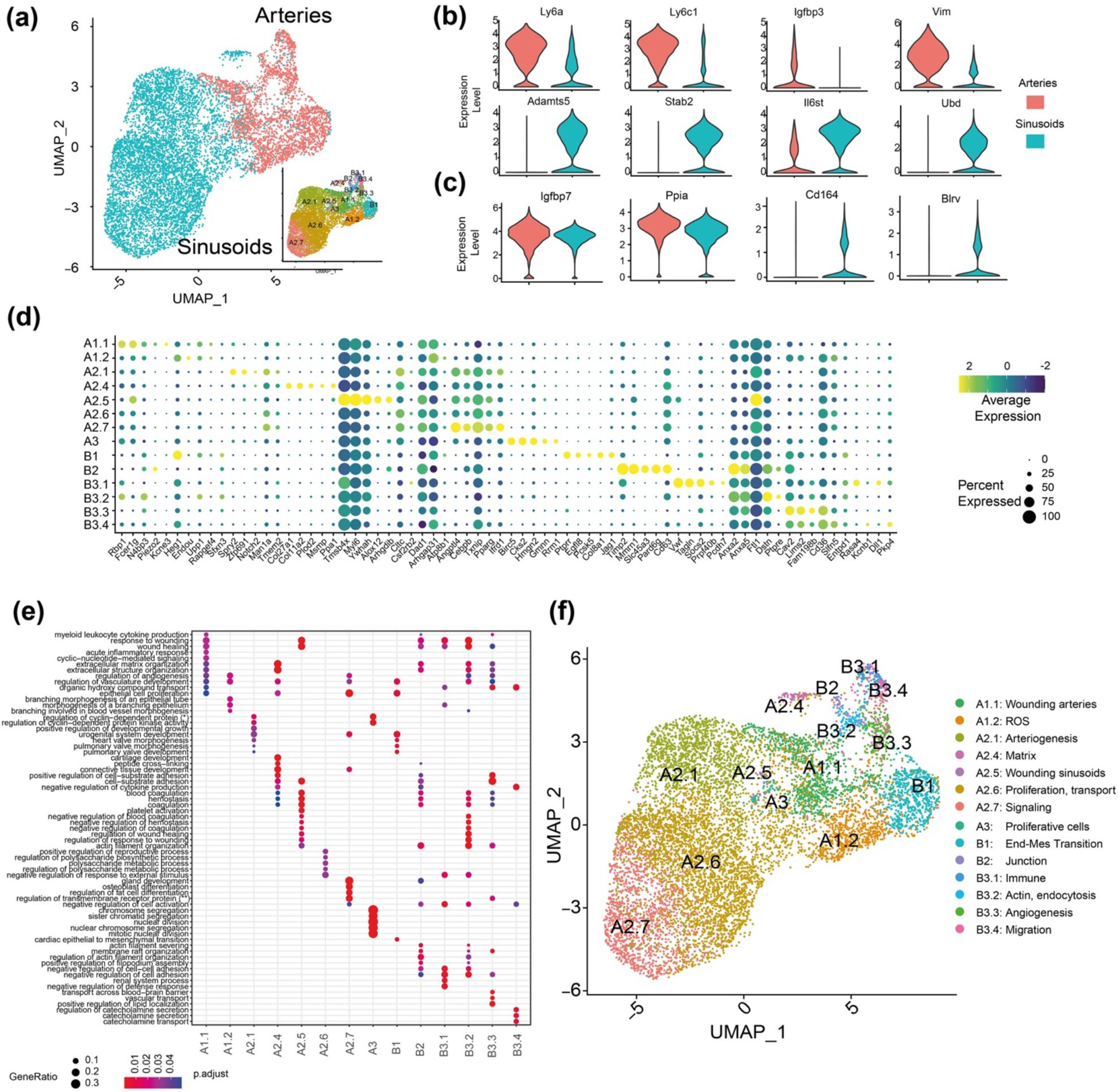
Deep characterization of the endothelial cell compartment in the BM. **(a)** UMAP representation of arteries (red) and sinusoids (blue) within the endothelial cell population. The right-bottom subpanel depicts the final endothelial clusters identified. **(b)** Violin plot of gene expression for known markers of arteries and sinusoids cell sub-types. **(c)** Violin plot of gene expression for new marker candidates separating arteries and sinusoids cell sub-types. **(d)** Dot plot of the top 5 markers for each endothelial subcluster. Dot size corresponds to the proportion of cells within the group expressing each gene, and dot color correspond to the average expression level. **(e)** Selected set of gene-sets derived from the gene-set analysis conducted with top 50 markers per cluster (see Methods). **(f)** Final clustering of the endothelial cell population and the labeling assigned based on marker genes and gene-set analysis.

Beyond characterizing arteries and sinusoids, we annotated their respective cell states using clusters markers based on genes **(Table S1, Fig. 4d)** and gene sets (derived from gene set enrichment analysis; **Table S3, Fig. 4e)**. Our final annotation is described in **Fig. 4f**. Gene sets related to vasculature development and remodeling were identified within the endothelial subclusters, confirming the identity of this cell population **(Table S3)**. We uncovered two subclusters (A1.1 and A2.5, respectively) within the arteries and sinusoids groups, which were enriched in gene sets involved in wounding. This finding is in agreement with the role of EC in pro- and anti-thrombotic processes(Yau et al., 2015). Gene sets involved in extracellular matrix assembly, cell adhesion, and migration processes were specifically enriched in arteries, in line with the importance of these biological processes for vascular morphogenesis(Davis and Senger, 2005). In relation to the structural support provided by arterial cells, subcluster B3.2 (actin, endocytosis) implicated in matrix remodeling was defined by the expression of *RhoC, Apln*, and *Anxa2*. Other arterial subclusters such as ROS and Immune (A1.2 and B3.1, respectively) include highly expressed gene sets involved in the regulation of reactive oxygen species metabolic process and cytokine-mediated signaling pathway. These findings are in line with the role of ECs in maintaining the REDOX balance and leukocyte regulation(Testa et al., 2016; Zhao et al., 2012). Sinusoidal-endothelial subclusters such as A2.1 and A2.6 showed enrichment in GO terms related to artery development and endothelial proliferation processes, two critical steps within the process of angiogenesis(Naito et al., 2021). Furthermore, the sinusoidal subcluster A2.7 expressed gene sets involved in ion transport and signaling-related signatures. This is in concordance with the need of EC to constantly sense and adapt to alterations in response to microenvironmental cues(March et al., 2009; Quillon et al., 2015). Of note, ion channels play a role in EC functions controlled by intracellular Ca^2+^ signals, such as the production and release of many vasoactive factors such as nitric oxide. In addition, these channels are involved in the regulation of the traffic of macromolecules, controlling intercellular permeability, EC proliferation, and angiogenesis(Nilius and Droogmans, 2001). Importantly, several markers that were found only with the dataset integration correspond to genes within the GO categories used to label the clusters, hence, revealing their important role in defining the function of these cell states. For example, in subcluster B2, genes such as, *Gja1, Tgfb3*, and *Ablim2* are involved in regulating cell junctions and cytoskeletal organization(Barrientos et al., 2007; Okamoto and Suzuki, 2017). Taken together, these results suggest a remarkable level of specialization of the bone marrow endothelial cells. The specificity of the distinct functional states indicates that the endothelial compartment is a more dynamic and flexible tissue with a richer intrinsic repertoire than previously anticipated.

### Deep characterization of the BM mesenchymal cell compartment

Applying the same robust clustering to mesenchymal stromal cells, we identified 11 subclusters and proceeded with the annotation **(Fig. 3c, Fig. S3, and Table S4-S6)**. Based on the expression of canonical markers, we first discriminated clusters between early mesenchymal (MSC), and cells already committed to the osteolineage (OLN-primed) **(Fig. 5a and Table S5)**. The high expression of *Cxcl12, LepR, Adipoq*, or *Vcam1*, among others, confirmed the identity of the early MSC group **(Fig. 5b left column)**; whereas *Bglap, Cd200, Alpl*, or *Col1a1* expression revealed the presence of osteolineage committed cells within the mesenchymal compartment **(Fig. 5b right column)**. Furthermore, we identified a number of previously unrecognized, differentially expressed genes between the MSC and OLN-primed clusters such as *Sbds* and *Itgb1* for MSCs and *Enpp1* and *Vkorc1*, for OLN-primed cell type population **(Fig. 5c)**. *Itgb1*, highly expressed in MSC is implicated in human chondrogenic differentiation of mesenchymal cells(Hamidouche et al., 2009). Among OLN-primed specific markers, *Enpp1* and *Vkorc1* have been shown to regulate bone development by regulating bone calcification(Hajjawi et al., 2014; Mackenzie et al., 2012; Price, 1985; Spohn et al., 2009).

**Figure 5.**
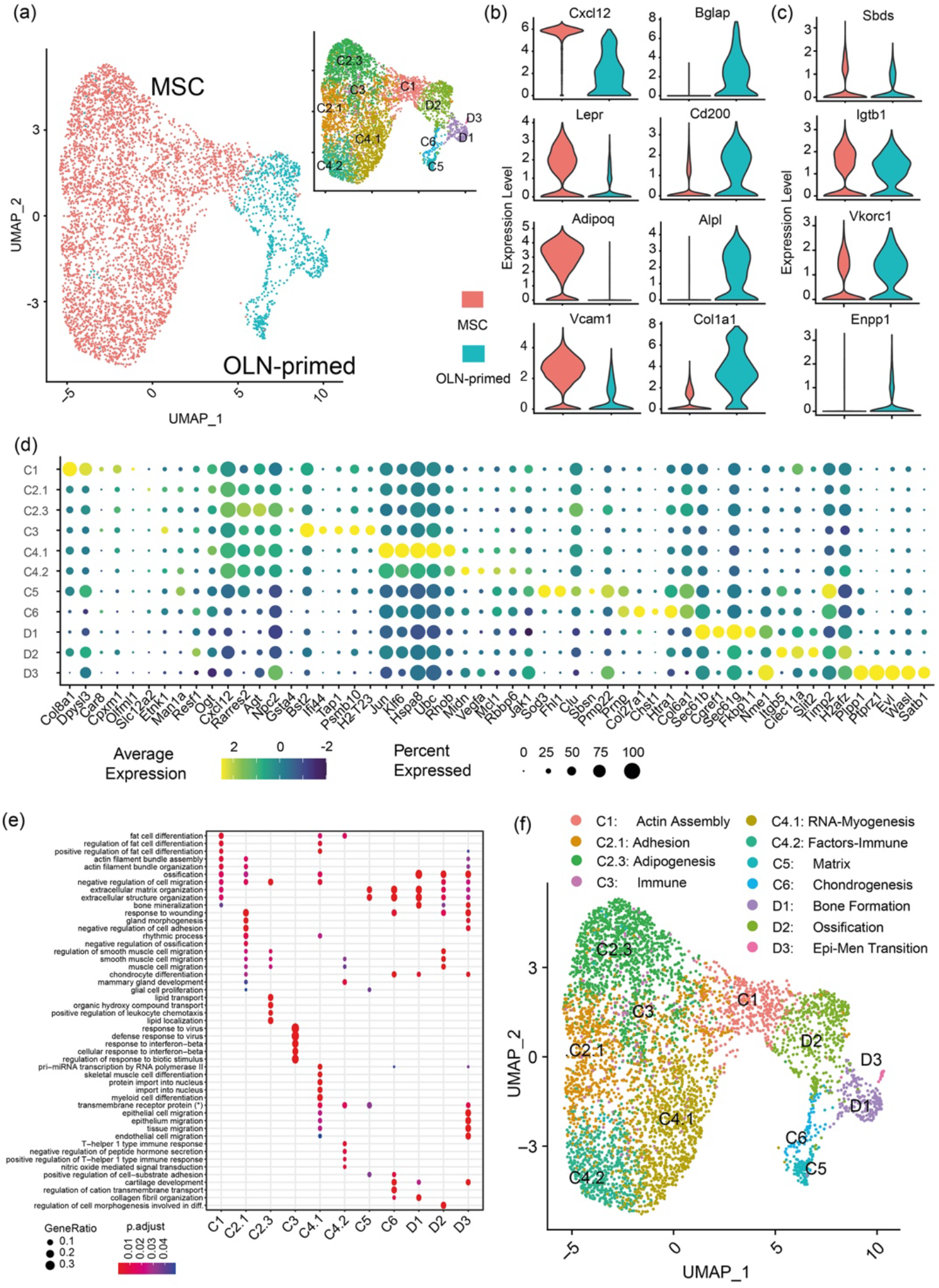
Deep characterization of the mesenchymal cell compartment in the BM. **(a)** UMAP representation of mesenchymal (red) and Osteolineage-primed (OLN-primed) (blue) within the mesenchymal compartment. The right-upper subpanel depicts the final mesenchymal clusters identified. **(b)** Violin plot of gene expression for known markers of Mesenchymal (red) and Osteolineage-primed (blue) cells. **(c)** Violin plot of gene expression for new marker candidates separating Mesenchymal (red) and Osteolineage-primed (blue) cells. **(d)** Dot plot of the top 5 markers for each mesenchymal subcluster. Dot size corresponds to the proportion of cells within the group expressing each gene, and dot color correspond to the average expression level. **(e)** Selected set of gene-sets derived from the gene-set analysis conducted with top 50 markers per cluster (see Methods). (*) *transmembrane receptor protein serine/threonine kinase signaling pathway*. **(f)** Final clustering of the mesenchymal cell population and the annotation based on marker genes and gene-set analysis.

Besides MSCs and OLN-primed MSCs, we identified additional subpopulations. Through the marker identification and gene set analysis of the 11 sub-clusters **(Fig. 5d,e and Table S4**,**6)** we were able to characterize and label each of the clusters as shown in **Fig. 5f**. GO terms such as adipogenesis, assembly and organization, immune response, cell migration or muscle differentiation were enriched in the C2.3, C2.1, C1, C3, C4.2 and C4.1 subsets respectively, confirming the identity of this MSC cell group. Furthermore, terms related to extracellular matrix, chondrocyte differentiation and bone development, including bone formation, ossification, or epithelial migration, among others, were identified in the OLN-primed subclusters (C5, C6, and D respectively), verifying the identity of these more mature cells (OLN-primed cells) within the mesenchymal stromal cells.

Taken together, these results demonstrate that the newly identified mesenchymal subpopulations could not be properly characterized without the multi-dataset integration and a novel clustering approach. Further, our data provide evidence of the heterogeneity of the mesenchymal compartment in the BM.

### Composition of the human endothelial and mesenchymal BM microenvironment

While our data reveal a previously unrecognized heterogeneity in the murine BM endothelial and mesenchymal compartments, information about the composition of the human microenvironment and how much of this heterogeneity is observed in humans remains unanswered. To address this issue, we performed scRNA-seq analysis in prospectively isolated EC (TO-PRO-3^-^, CD45^-^, CD235^-^, Lin^-^, CD31^+^ and CD9^+^(Barreiro et al., 2005; Kenswil et al., 2018)) and MSC-OLN (TO-PRO-3^-^, CD45^-^, CD235^-^,Lin^-^,CD271^+^(Ghazanfari et al., 2016; Hashemi et al., 1991; Quirici et al., 2002), CD146^+/-^) **(Fig. S5a)** from iliac crest BM aspirates from four healthy young adults (20-30 years of age) (**Fig. 6a**). As described in **Fig. 6b**, we added additional filtering steps in the bioinformatic analysis to identify the two populations of interest, EC and MSC. As an additional quality control, we estimated the contribution of each human sample to EC and MSC subsets and cell cycle status (**Fig. S5b**,**c)**. The EC (907 and 658 cells, clusters 1 and 6 respectively) (**Fig. 6c and Fig. S5b middle panel**) identity was confirmed based on the expression of canonical endothelial markers such as *PECAM1* (coding for *CD31*), *CD9, ICAM2, VLC*, and *ITGB1* (**Fig. 6d and Table S7**). In addition, examining functional pathways in clusters 1 and 6 revealed enrichment in GO terms associated with blood coagulation and hemostasis **(Table S8)**. The MSC identity (249 cells, cluster 11, **Fig. 6c and Fig. S5b middle panel**) was confirmed by the expression of the mesenchymal specific genes (*CXCL12 and LEPR*) and the OLN-Primed specific genes *ANGPT1, COL1A1, and VCAM1*, among others (**Fig. 6e and Table S7**). Further, enrichment in functions associated with extracellular matrix organization and response to the mechanical stimulus was demonstrated in osteolineage cells **(Table S8)**. In summary, the generated human data suggests that single-cell RNA sequencing from iliac crest aspirates can aid in describing the complexity of the human BM microenvironment. Nevertheless, the limited number of EC and MSC, as well as the presence of contaminating populations did not allow a fine-grained clustering as the one performed in the mouse data.

**Figure 6.**
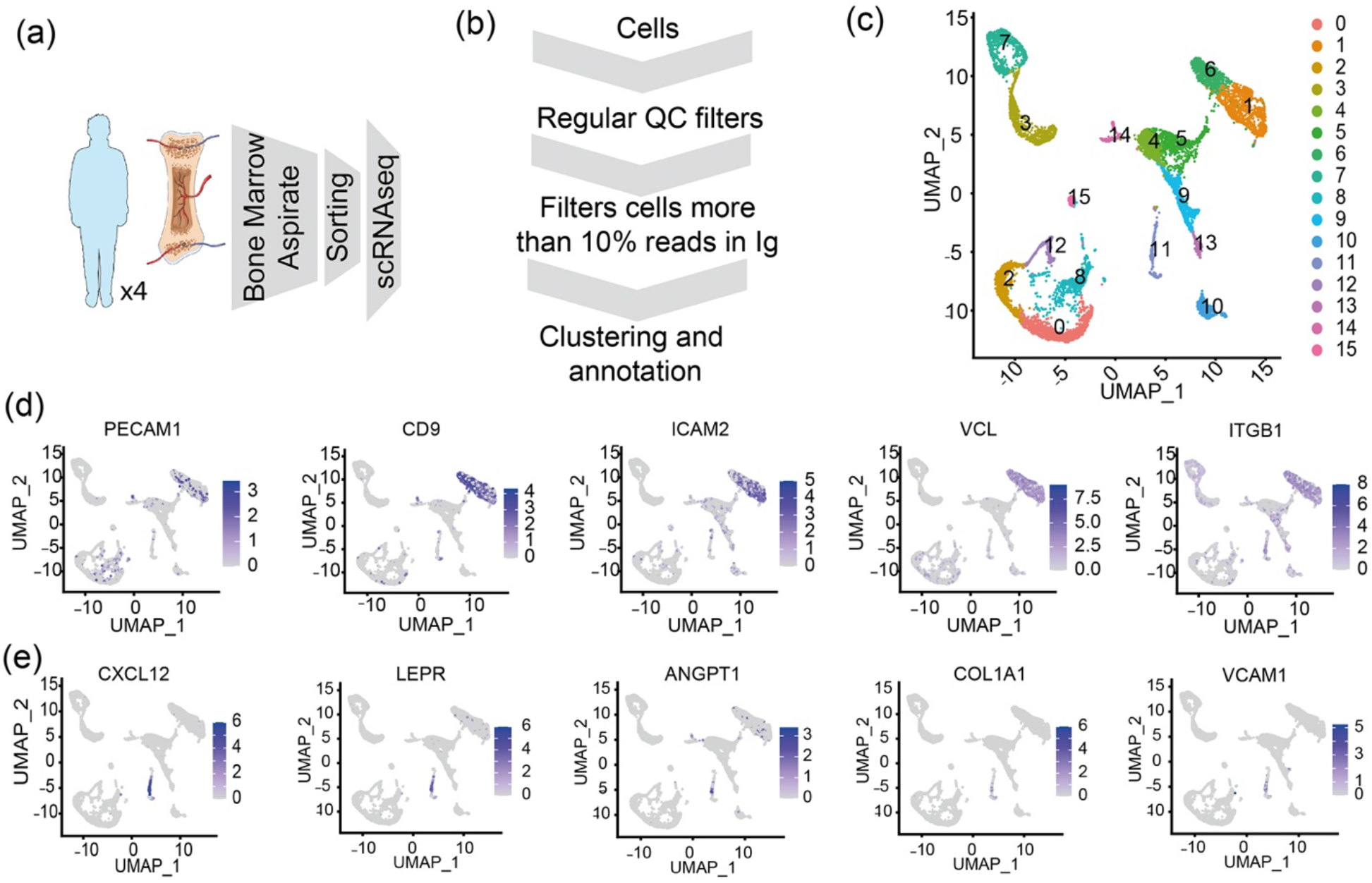
Composition of the human endothelial and mesenchymal BM microenvironment. **(a)** Experimental design for the human BMM characterization. **(b)** Scheme of customized bioinformatics pipeline filtering the cells with a large number of Immunoglobulin genes. **(c)** UMAP visualization of color-coded clustering of the human BM microenvironment after filtering cells. **(d**,**e)** Expression of representative markers for endothelial population (d) and mesenchymal-osteolineage cells (e) using an UMAP representation.

### Substantial conservation of the EC and MSC population in the human BM microenvironment

Based on the limitations of the human data, we next investigated to what extent the knowledge uncovered in mouse could be applied to identify subpopulations/cell states in the human BM microenvironment. As a first step, we used single-cell mouse data to annotate the human cells using SingleR(Aran et al., 2019), separately for EC **(Fig. S6a**,**b)** and MSC **(Fig. S6c**,**d)**. We observed that MVG identified in mouse allowed us to separate the cells into clusters for both human EC and MSC **(Fig. S6a**,**b**,**c**,**d)**. As a result, this analysis suggests that part of the biological mechanisms defining the BM microenvironment may be shared between species.

Therefore, we decided to investigate the enrichment of conserved features (genes) between mouse and human; therefore an enrichment score (ES) was computed for each cluster for EC and MSC separately (see Methods) **(Fig. 7a and Table S8-S9)**. In the case of endothelial cells the enrichment score was up to two-fold **(Fig. 7a)**: wounding (A1.1) with 2.15-ES, the junction (B2) 2.26-ES, arteriogenesis (A2.1) 2.02-ES and Signaling (A2.7) 2.5-ES). Importantly, for some of the subclusters as sinusoidal signaling (A2.7) and the arterial of angiogenesis (B3.3), these shared genes are critical for defining each of those EC functional states **(Table S9)**. Both human and mouse ECs express *DDIT4, JUN, CITED2, GADD45G, DUSP1, FOS*, and *CLDN5*, which are part of a wide array of transcription factors, growth factors, and signaling pathways that have been described to regulate the maintenance of vascular homeostasis under physiological conditions(Echavarria and Hussain, 2013; Escudero-Esparza et al., 2012; Jia et al., 2016). Similarly, ECs subclusters involved in angiogenesis in both species shared the expression of *RGCC, GATA2, KLF2*, and *CAV2* genes, which are known to be implicated in angiogenic related processes(Lee et al., 2006; Linnemann et al., 2011).

**Figure 7.**
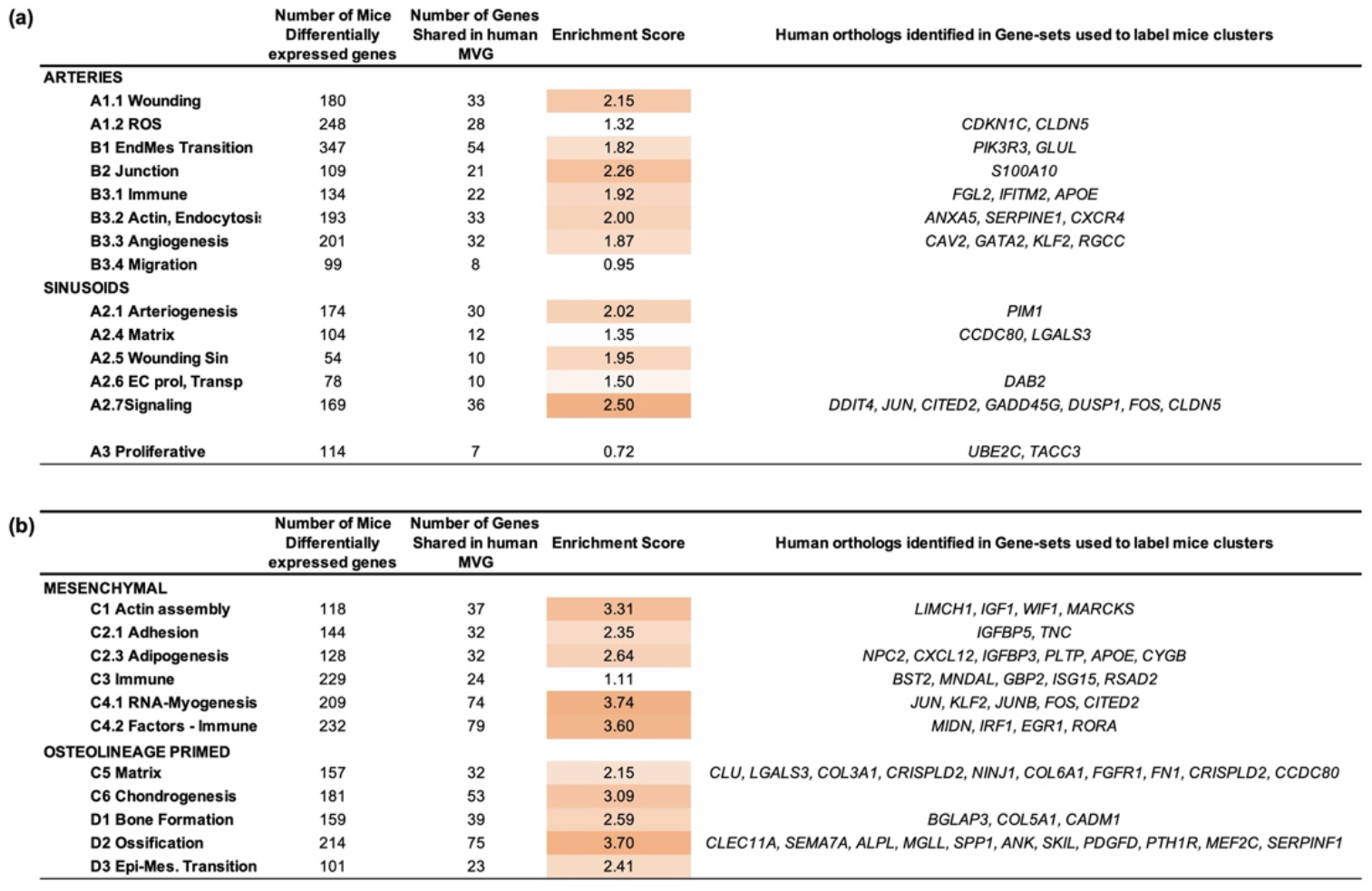
Conservation analysis of the EC and MSC population in the human BM microenvironment. **(a,b)** Quantification of the conservation for EC (a) and MSC-OLN (b) cells for each cluster. The enrichment of those genes that are cluster markers in mouse and observed in most variable genes (MVG) of EC and MSC human cells respectively. The right column shows, among the genes identified in human, those that are part of the gene-sets used to label the cluster.

In the case of mesenchymal cells, we identified >3.5-ES for three subclusters, such as RNA-Myogenesis (C4.1), Factors-Immune (C4.2), and ossification (D2). Importantly, some of the shared genes correspond to genes that allowed the subcluster labelling through GO categories **(Fig. 7b and table S8 and S10)**. Genes such as *CXCL12, APOE*, or *IGFBP3* are associated with cell migration and lipid transport pathways among others(Amable et al., 2014; Robert et al., 2020), and characterized the murine adipogenesis subcluster (C2.3). Other genes, such as *IGFBP5*, are involved in actin filament assembly and organization(Sureshbabu et al., 2012) and defined the cell adhesion subcluster (C2.1). *IFIT3, MIDN, and ILR1* belong to pathways associated to interferon regulation or autoimmunity(Guo et al., 2017; Kim et al., 2020) and were identified in the immune response subclusters (C3 and C4.2). Additionally, the expression of *COL5A1* and *CADM1* genes, previously related to collagen fibril organization and bone mineralization processes(Kahai et al., 2004; Nakamura et al., 2017), defined the bone formation subcluster (D1). Moreover, genes such as *SPP1* or *CLEC11A*, which are related to osteoblasts function and mineralization(Huang et al., 2007; Shen et al., 2021; Yue et al., 2016) defined the mouse OLN-primed MSCs subcluster associated to ossification (D2) and are also expressed in human MSC-OLN cluster. Altogether, this data indicates the conservation of the osteogenic microenvironment between both species.

Together, our analysis suggests that deep characterization of cellular states in mice can be used to infer conserved features in the human BM microenvironment despite a low level of conservation in the actual transcriptional profile of EC and higher in the case of MSC. Importantly, our data reveal a substantial degree of conservation regarding the complexity and heterogeneity of the EC and MSC compartment in the BM between mouse and human. This suggests that the layers of microenvironmental regulation of hematopoiesis and the identified plasticity in mice may also be shared between species.

## DISCUSSION

Our study dissects the intrinsic organization and the heterogeneity within the endothelial (EC) and mesenchymal cell populations (MSC) governing the BM microenvironment. This was accomplished through customized bioinformatics integration of multiple datasets along with the inclusion of over 50.000 murine bone marrow stromal cells. We were able to identify new subsets of MSC and EC but, but more importantly, to define new molecular markers for the identification of highly specialized subpopulations of cells in the BM microenvironment. Pathway enrichment analysis unveiled multiple, potentially transient cell states defined by differential gene expression and the enrichment of specific functional characteristics. Importantly, EC subsets were characterized by enrichment in pathways known to be essential for endothelial homeostasis maintenance, demonstrating a high degree of specialization in the endothelium. Similarly, multiple transient cell states in the MSC compartment were defined and characterized by their differentiation capacity. Importantly, our deep deconvolution of the heterogenous mesenchymal and endothelial compartments became feasible only by integrating multiple datasets. Of note, our analysis showcases that a research paradigm aiming for the generation of a detailed comprehensive molecular atlas of an organism requires both multi-omic data and computational integration. Here, we have relied on what is referred to as unpaired unimodal (scRNA-seq) data. Clearly, a natural next step is to use and further develop new computational tools that permit the integration(Argelaguet et al., 2021) of unpaired multi-omics datasets such as scRNA-seq, scATAC-seq, and other data modalities. Recent technological developments enable several multiple omics recorded from the same cell, i.e. paired data, which leverages our ability to dissect and molecularly characterize the intrinsic organization of the bone marrow niche environment. Advances in computational biology have started to develop such tools(Ashhurst et al., 2021; Hao et al., 2021; Martinez-de-Morentin et al., 2021; Wu et al., n.d.). Moreover, some validation experiments, such as the use of fluorescent reporters, targeting niche-associated genes and functional studies will allow us to confirm the identified new molecular markers based on differential gene expression and also the related annotated pathways.

While our study did not directly address the influence between stromal cells in the hematopoietic stem cell niche and the HSCs, the deep resolution of our study allows for some inferences to be made. Of note, we detected the expression of vascular endothelial growth factor-C (*Vegf-c*) in mouse endothelial and mesenchymal cells **(Table S1, S4, S5, S7, and S10)**. *Vegf-c* has recently been implicated in the maintenance of the perivascular niche and the recovery of hematopoiesis upon injury(Fang et al., 2020). *Vegf-c* is specifically expressed in the endothelial B1 and mesenchymal C1 subpopulations, suggesting an important role of these specialized endothelial and mesenchymal cells in the preservation of the integrity of the perivascular niche. In addition, Apelin+ (Apln) endothelial cells have been recently implicated in HSC maintenance and regeneration upon injury(Q. Chen et al., 2019). Importantly, two endothelial subclusters (B3.2-actin endocytosis and A1.1-wounding arteries) demonstrated expression of Apln, suggesting that these EC states represent specific sources of hematopoietic support and vascular regeneration upon injury. Osteolectin+ LepR+ - mechanosensitive peri-arteriolar mesenchymal cells with osteogenic potential - are implicated in lymphoid, but not HSC maintenance(Shen et al., 2021). Importantly, osteolectin expression defines murine cluster D2 (ossification) and shows conservation in human MSCs, suggesting the preservation of a specialized lymphoid niche between species.

Detailed characterization of the human BM niche has not yet been addressed. Approaches undertaken in a mouse system cannot readily be transferred to the human system. Furthermore, differences in sample processing can also impact the results. In that sense, our results, despite the low number of cells, may represent the first dataset that includes scRNAseq from the human endothelial and mesenchymal BM microenvironment. While we were able to identify mesenchymal and endothelial cells based on canonical markers shared with mice(Aoki et al., 2021; Baryawno et al., 2019; Kalucka et al., 2020; Leimkühler et al., 2021; Matthews et al., 2020; Severe et al., 2019; Stumpf et al., 2020; Tikhonova et al., 2019; Wang et al., 2020; Xie et al., 2021), our human scRNAseq did not possess enough resolution to elucidate the heterogeneity of the human BM stroma to the same level as with the mouse data. Based on the extensive knowledge generated in the mouse, we therefore focused on characterizing how much of the information and targets from the mouse can be of interest in human characterization. This analysis allowed us to identify the expression of the human orthologs to the murine cluster-defining genes with different degrees of enrichment in the endothelium and mesenchyme. Moreover, some of these shared genes in mice and human stromal cells corresponded to the GO-defining genes of the different clusters identified in the mouse. These findings suggest a meaningful degree of conservation regarding the cellular states that define the stromal microenvironment in mouse and human. Although additional studies and improved processing of human samples will be required for deep characterization of the human BM microenvironment, these preliminary results validate our integrative cross-species approach.

As an example of the added value, the current study identifies candidates of relevance in the study of BM related diseases. *Sbds*, a ribosome maturation protein associated with the Shwachman-Diamond syndrome, represent a previously unrecognized marker of immature MSCs based on the dataset integration. Sbds deficiency has been implicated in ossification defects and metabolic changes in HSPCs(Raaijmakers et al., 2010; Zambetti et al., 2016), potentially contributing to myelodysplasia and AML onset in Shwachman-Diamond syndrome patients. Future studies will help to improve the understanding of these new candidates and the pathogenesis associated. On a broader note, deep molecular analysis of the BM microenvironment set the stage for computational disease modelling(Tegnér et al., 2009) and

Taken together, our study provides a deeper understanding of the composition and specialization of the BM microenvironment and point towards a substantial degree of conservation between species. Moreover, we demonstrate the usefulness of the multi-dataset integration and the customized clustering approach used in our study to improve the resolution of complex tissues and organs. This approach promises to aid in the construction of cell atlases by reducing the resources associated with sequencing that a single lab will need to invest in order to obtain meaningful depth in single-cell analysis while reducing the biases that may arise from data generated from a single laboratory or platform.

Future studies integrating genome, transcriptome, epigenome, proteome, and anatomical positioning together with functional assays to correlate descriptive phenotypes with functional data will help fully resolve the composition, regulation, and connectivity in the BM microenvironment in health and disease.

## METHODS

### Mouse and human samples

Female C57BL/6J mice (CD45.2, Jackson Laboratory #000664) at age 20 weeks were used for scRNA-seq experiments. All animal experiments were performed in accordance with national and institutional guidelines and procedures were approved by the Ethical Committee for Animal Testing of the University of Navarra.

The human sample collection and research conducted in this study were approved by the Research Ethics Committee of the University of Navarra. All the protocols used in this study were in strict compliance with the legal and ethical regulations. After informed consent, a total volume of approximately 60 mL bone marrow was obtained by aspiration from the posterior iliac crest from healthy young volunteers (20-30 years of age).

### Isolation and FACS sorting of murine bone marrow microenvironment cells

Mice (x6) were euthanized via CO2 asphyxiation. Bones from humerus, radius, iliac crests, femurs and tibia were harvested in PBS 1X containing 2 % FBS and 2 mM EDTA (modPBS). All steps were performed on ice to preserve cell viability and RNA integrity. Muscles and soft tissue were thoroughly removed from the bones and BM cells were obtained by crushing in modPBS. Cells were then filtered through a 70 μm cell strainer and red blood lysed with ACK buffer (NH4Cl 150 mM, KHCO3 10 mM, and Na2EDTA 0,1 mM) for 10 minutes at room temperature (RT). Remaining calcified bone fragments were collected on a 50 mL conical tube and digested with the appropriate volume of PBS with 0.3% collagenase I and dispase (5 U/ml) during 15 min at 37ºC and shaking at 200 rpm. FBS representing 10% of the digestion volume was added to stop the collagenase digestion. After digestion, the calcified and crushed fractions were filtered through a 70 μm filter into a collection tube and pooled into one sample. Cells were subsequently stained for 20 minutes on ice first in the appropriate volume of modified PBS 1X (3 ml/mouse) with 160 ul/mouse of biotinylated lineage cocktail (Mac1, CD3, Gr1, B220 and TER119) followed by incubation with streptavidin magnetics microbeads (100 μl/mouse). Negative selection was performed using Miltenyi LD columns according to manufacturers’ protocol.

After selection, the sample was stained with the following combination of conjugated antibodies at a concentration of 1/200: APC-Cy7 labeled streptavidin, BV510 labeled anti-CD45, APC labeled anti-CD45 and APC labeled anti-Ter119. Samples were then stained with 0.05 μM of Vybrant dye orange (VDO) at 37ºC for 30 minutes to label living cells. Annexin V was also added, in combination with 7AAD to discard apoptotic and dead cells from the sample respectively. For annexin V staining, cells were stained with 1 μl/mouse of Annexin V-FITC on an appropriate volume of 1X Annexin V binding buffer in the dark for 15 min at RT. Samples were then resuspended in 1X Annexin V buffer and 5 μl of 7AAD dye (up to 1×10^6^ cells) were added. BM non-hematopoietic cells were FACS sorted using BD FACSAria II sorter collected in PBS 1X supplemented with 0.05% UltraPure BSA and cell viability was assessed using Nexcelom Cellometer. Data were analyzed by FACSDiva (BD) or FlowJo (version 10.7.1) software.

### Isolation and FACS sorting of human bone marrow endothelial and mesenchymal-osteolineage cells

All sample processing steps were performed on ice to preserve cell viability and RNA integrity. A total volume of approximately 60 ml of bone marrow was obtained by aspiration from the posterior iliac crest. Red blood cells were lysed twice with 45 ml of ACK buffer per 5 ml of human sample during 15 minutes at RT with rotation. Sample was then filtered through a 70 μm cell strainer, centrifuged, and stained for 30 min on ice with the following combination of conjugated antibodies at a concentration of 1/100 except anti-Lin (3ul/test-test 25×10^6^cells): BV510 labeled anti-Lin (including CD3, CD10, CD19, CD45 and CD64), BV421 labeled anti-CD235, BV421 labeled anti-CD45, FITC labeled anti-CD31, APC-Cy7 labeled anti-CD9, PE labeled anti-CD146 and PerCP-Cy5.5 labeled anti-CD271. Dead cells and debris were firstly excluded by FSC, SSC and adding 10 μl of TO-PRO-3. BM niche populations were prospectively isolated based on the following immunophenotype: ECs: TO-PRO-3^-^/Lin^-^/CD45^-^/CD235^-^/CD9^+^/CD31^+^ and MSCs: TO-PRO-3^-^ /Lin^-^/CD45^-^/CD235^-^/CD31^-^/CD271^+^/CD146^+/-^. FACS sorting was performed on a BD FACSAria II sorter, sorted BM niche cells were collected in PBS 1x and 0.05% UltraPure BSA and cell viability was determined using Nexcelom Cellometer. Data were analyzed by FACSDiva (BD) or FlowJo (version 10.7.1) software.

### Profiling by Single-cell RNA-sequencing (scRNA-seq)

scRNA-seq was performed using the Single Cell 3’ Reagent Kits v3.1 (10X Genomics) according to the manufacturer’s instructions. For human samples, endothelial and mesenchymal cells were pooled before scRNA-seq was performed. Approximately 15,000 cells were loaded at a concentration of 1,000 cells/µL on a Chromium Controller instrument (10X Genomics) to generate single-cell gel bead-in-emulsions (GEMs). In this step, each cell was encapsulated with primers containing a fixed Illumina Read 1 sequence, followed by a cell-identifying 16 bp 10X barcode, a 10 bp Unique Molecular Identifier (UMI) and a poly-dT sequence. A subsequent reverse transcription yielded full-length, barcoded cDNA. This cDNA was then released from the GEMs, PCR-amplified and purified with magnetic beads (SPRIselect, Beckman Coulter). Enzymatic Fragmentation and Size Selection was used to optimize cDNA size prior to library construction. Illumina adaptor sequences were added, and the resulting library was amplified via end repair, A-tailing, adaptor ligation and PCR. Libraries quality control and quantification was performed using Qubit 3.0 Fluorometer (Life Technologies) and Agilent’s 4200 TapeStation System (Agilent), respectively. Sequencing was performed in a NextSeq500 (Illumina) (Read 1: 26 cycles, i7 Index: 8 cycles, Read 2: 49 cycles) at an average depth of 60,000 reads/cell in mice and 30,000 reads/cell in human.

### Single-cell RNA-seq Analysis of mouse samples

See extended details and code in the following Github: https://github.com/TranslationalBioinformaticsUnit/BMN_characterization.

#### Sample selection

Sample GSM3674224, GSM3674225, GSM3674226, GSM3674227, GSM3674228, GSM3674229 from GSE128423 by Baryawno, sample GSM2915575, GSM2915576, GSM2915577 from GSE108891 by Tikhonova and one in-house mouse bone marrow niche sample was included in this analysis.

#### Filtering

the single cell analysis of mice samples analysis was performed using R (version 4.0.3, 3.6.3) and Seurat (version 4.0.0, 3.2.3)(Stuart et al., 2019). Three bone marrow niche samples were filtered individually based on the 10^th^ and 90^th^ quantile of number of features and counts. Cells with more than 5% mitochondrial genes were also removed. Each dataset was normalized using SCTransform(Hafemeister and Satija, 2019) separately.

#### Pairwise integration and selection of the target population

In-house dataset and Baryawno were integrated with Tikhonova separately using Seurat (version 3.2.3.). Using as a reference the annotation from Tikhonova dataset, cells that aligned with LEPR+ cells and VE-Cad+ cells were annotated as MSC and EC respectively. MSC-like cells and EC-like cells from different datasets were normalized again and integrated using Seurat without further filtering.

#### Clustering

After filtering and quality control, a divide-and-conquer strategy was applied to the clustering of mouse ECs and MSCs separately. Firstly, following integration, dimension reduction with principal component analysis (PCA), data visualization with Uniform Manifold Approximation and Projection (UMAP), computation of K-nearest neighbors and clustering using resolution of 1 were performed as a reference of high-resolution limit. Secondly, IKAP(Y. C. Chen et al., 2019) was applied to each integrated dataset as “level 1” clustering (Fig. 2,b). Each cluster from level 1 was then compare with the high-resolution reference. The cluster from level 1 was further divided using IKAP to level 2 if the cluster were far from cluster limit. The process would be repeated until at least one cluster reach cluster limit.

#### Cluster evaluation

To evaluate the stability of these clusters, a bootstrapping strategy was adopted(Tasic et al., 2016b). The strategy was conducted in a pairwise manner with basic steps as follow:

1. Select two clusters, randomly split the clusters to five equal groups and use one group of cells (20%) as testing dataset.
2. Identify the set of differentially expressed genes (DEGs) between the 2 clusters using the Wilconxon Rank Sum test.
3. Train a random forest classifier with 20 genes selecting the top 10 DEGs from each cluster based on average log2 fold change.
4. Applied the classifier to the 20% testing dataset.
5. Repeat step1-4 for five times for different groups such that each cell in these two clusters was classified once.
6. Repeat step1-5 nine more times.
7. Repeat step1-6 for all cluster pairs

There are three types of results that can be summarized from this bootstrapping strategy:

1. Dominant cluster identification: A dominant cluster is the cluster to which the cell is assigned for more than half the runs.
2. Number of Correct Dominance Assignment (#Correct): The sum of times the dominant cluster matches the original cluster (true positive) for a cell across all cluster pair comparisons (Fig. S2e).
3. Recall Per Cell: The proportion of correct assignment (positive result) to its original cluster in all runs from all comparisons (Fig. S2d).

Clusters where more than 50% of the cells has been “incorrectly” assigned robustly at least once to another dominant cluster will be considered unstable (Fig. S2, cluster A2.2). The cells of such cluster will be assigned to other clusters (see dominant plots, **Fig. S7** as an example) using – to that end - a random forest classifier as described before.

#### Gene Set analysis

After clustering, DEGs of each cluster were identified using a Wilconxon Rank Sum test. For each cluster, the relevant gene sets were identified using the top 50 DEGs using clusterProfiler (version 3.18.1)(Yu et al., 2012).

### Added value analysis

#### Added value 1

Comparing the DEGs defined by individual dataset and integrated dataset. The individual datasets were normalized with SCTransform and DEGs were identified within top 3000 most variable genes using a Wilconxon Rank Sum test. For integrated dataset, DEGs were identified within top 3000 most variable genes from integrated assay using a Wilconxon Rank Sum test. False negative and false positive rate were calculated by comparing the DEGs identified by integrated dataset and individual dataset (Fig. 3e,f).

#### Added value 2

Cluster stability evaluation for individual dataset. To understand if the clusters identified from three datasets can remain stable within a single dataset or not, the bootstrapping strategy was applied to each dataset with the annotation identified by the integrated dataset.

#### Added value 3

Comparing cluster identified by a single dataset. To further understand if the clusters can be identified by one dataset only, Baryawno dataset was used as an example considering its large cell populations. The same pipeline from normalization to bootstrapping was applied to this dataset and the clusters identified from this single dataset was compared with the clusters identified by three datasets using Jaccard index.

### Single-cell RNA-seq Analysis of Human samples

#### Preprocessing of sequencing data

Preprocessing of single-cell RNA-seq data for each in house sample were conducted by CellRanger count from Cell Ranger (version 6.0.1) using reference genome GRCh38.

#### Sample filtering

The single cell analysis of human analysis was performed as described in before except for human cells with more than 10% mitochondrial genes were also removed. Because the exploratory data analysis revealed potential contamination of B cells, we applied an additional filter: cells with more than 10% reads mapped to immunoglobin genes were excluded from downstream analysis.

#### Integration

After filtering, each sample was normalized using SCTransform and integrated using 3000 most variable features using Seurat. Following integration, dimension reduction with PCA, data visualization with UMAP, computation of K-nearest neighbors with 20 dimensions and clustering using resolution of 0.4 were performed.

#### Select EC and MSC

(without further integration): Clusters were annotated based on biological insights on markers. Cluster 11 were identified as *mesenchymal* and cluster 1, 6 were identified as *endothelial*. During the exploratory analysis, human EC cells were subclustered at resolution 0.4 and one of the clusters identified was further filtered for the downstream analysis because the cells in the cluster were not expressing EC marker genes. Several outliers from human MSC cells were also removed.

#### Compare human MVGs with mouse DEGs: 3000 MVGs for human EC and MSC were identified using “RNA” assay and these genes were scaled in “integrated” assay resulting in 932

human EC specific MVGs and 976 human MSC specific MVGs. The MVGs from human EC or MSC were compared with DEGs from each mouse cluster. The enrichment score for a given cluster *i* was defined as the ratio between “*the number of genes shared between human MVGs and mouse DEGs from cluster i*” and “*the expected number of genes*”, where the later was computed as follows:

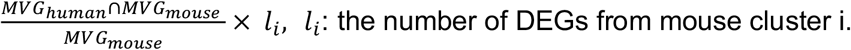

### SingleR analysis between mouse and human

To annotate human cells using mouse clusters as reference the singleR tool (version 1.4.1)(Aran et al., 2019) was utilized for ECs and MSCs separately. Cell type specific MVGs with expression values in “integrated” assay from Seurat object were used for this analysis.

## FIGURES

**<u>Figure 1. Overview of the paper.</u>**Graphical brief description of the paper.

**<u>Figure 2. Data integration and high-resolution clustering strategy.</u>(a)** Integrated analysis of the bone marrow niche datasets (two publicly available, Tikhonova et al. and Baryawno et al., and one generated in our lab, in-house) separately for two well-defined populations (endothelial and mesenchymal cells). Tikhonova et al. data set are used as a reference considering their separated cell profiling strategy for COL2.3+, LEPR+ and VE-Cad+ populations. In the top row, a UMAP projection is depicted for each single-cell RNA dataset. In the left-lower (“In-house + Tikhonova”) and in the right-lower (“Baryawno + Tikhonova”) data sets are integrated to identify endothelial and mesenchymal populations. (b) Clustering strategy: the analysis of endothelial cells as an example. An upper limit to the cluster is set for the clustering (left panel) using Louvain high-resolution clustering. Then, an iterative divide-and-conquer strategy identifies the optimal level of clusters at different levels: Level 1 (second panel from the left), Level 2 (third panel), and Level 3 (fourth panel). (c), (d), (e) The robustness analysis for sub-clustering B3 (from Level 2 to Level 3). Specifically: (c) sub-clusters identified, (d) the fraction of assignments to its original cluster using a random-forest + bootstrapping strategy and (e) summary of the results (d) per cluster, #correct indicates the times a cluster is a dominant cluster (see Methods) for a cell inside in all pairwise comparisons (see Fig.S2 for the sub-clustering analysis of A1 and A2).

**<u>Figure 3. Quality control and added value of the clustering analysis.</u>(a)** Representation of the final endothelial clusters. (b) Left panel: proportion of cells per dataset in each cluster in the final Endothelial clustering analysis. Right panel: proportion of cells per cell cycle stage using Seurat in each cluster in the final Endothelial clustering analysis. (c,d) Similar as (a,b) for MSC. (e,f) The added value of the integrated approach. Upper panel: every cell depicts the % of markers identified per cluster using only one dataset when compared with the markers identified in the total dataset. Middle panel: every cell depicts the % of False Negatives. Bottom panel: every cell depicts the % of False Positives when comparing the analysis conducted within each dataset with the integrated analysis (considered as the correct result). (e) and (f) are respectively associated with endothelial and mesenchymal cells. (g,h) Robustness of the cluster characterization using only cells from a single dataset but maintaining the same cluster structure. (g) and (h) are respectively associated with endothelial and mesenchymal cells.

**<u>Figure 4. Deep characterization of the endothelial cell compartment in the BM.</u> (a)** UMAP representation of arteries (red) and sinusoids (blue) within the endothelial cell population. The right-bottom subpanel depicts the final endothelial clusters identified. (b) Violin plot of gene expression for known markers of arteries and sinusoids cell sub-types. (c) Violin plot of gene expression for new marker candidates separating arteries and sinusoids cell sub-types. (d) Dot plot of the top 5 markers for each endothelial subcluster. Dot size corresponds to the proportion of cells within the group expressing each gene, and dot color correspond to the average expression level. (e) Selected set of gene-sets derived from the gene-set analysis conducted with top 50 markers per cluster (see Methods). (f) Final clustering of the endothelial cell population and the labeling assigned based on marker genes and gene-set analysis.

**<u>Figure 5. Deep characterization of the mesenchymal cell compartment in the BM.</u> (a)** UMAP representation of mesenchymal (red) and Osteolineage-primed (OLN-primed) (blue) within the mesenchymal compartment. The right-upper subpanel depicts the final mesenchymal clusters identified. (b) Violin plot of gene expression for known markers of Mesenchymal (red) and Osteolineage-primed (blue) cells. (c) Violin plot of gene expression for new marker candidates separating Mesenchymal (red) and Osteolineage-primed (blue) cells. (d) Dot plot of the top 5 markers for each mesenchymal subcluster. Dot size corresponds to the proportion of cells within the group expressing each gene, and dot color correspond to the average expression level. (e) Selected set of gene-sets derived from the gene-set analysis conducted with top 50 markers per cluster (see Methods). (*) transmembrane receptor protein serine/threonine kinase signaling pathway. (f) Final clustering of the mesenchymal cell population and the annotation based on marker genes and gene-set analysis.

**<u>Figure 6. Composition of the human endothelial and mesenchymal BM microenvironment.</u> (a)** Experimental design for the human BMM characterization. (b) Scheme of customized bioinformatics pipeline filtering the cells with a large number of Immunoglobulin genes. (c) UMAP visualization of color-coded clustering of the human BM microenvironment after filtering cells. (d,e) Expression of representative markers for endothelial population (d) and mesenchymal- osteolineage cells (e) using an UMAP representation.

**<u>Figure 7. Conservation analysis of the EC and MSC population in the human BM microenvironment.</u> (a,b)** Quantification of the conservation for EC (a) and MSC-OLN (b) cells for each cluster. The enrichment of those genes that are cluster markers in mouse and observed in most variable genes (MVG) of EC and MSC human cells respectively. The right column shows, among the genes identified in human, those that are part of the gene-sets used to label the cluster.

## Abbreviations

(HSC): Hematopoietic stem cell
(HSPC): hematopoietic stem and progenitor cells
(scRNA-seq): Single-cell RNA sequencing
(BM): Bone marrow
(EC): endothelial cells
(MSC): mesenchymal cells
(OLN-primed): osteolineage
(MVG): most variable genes

## ACKNOWLEDGMENTS

We would like to thank the staff of the flow cytometry core, the advances genomic lab and the animal facility at CIMA Universidad de Navarra for their invaluable technical and intellectual assistance. We are particularly grateful to the healthy volunteers who donated bone marrow tissue for this study. We also acknowledge Ali O. Balubaid’s help in writing.

This research was funded by grants from The Spanish Government, through project PID2019-111192GA-I00 (MICINN) to DGC. Instituto de Salud Carlos III (ISCIII) and co-financed by FEDER: PI16/02024, PI17/00701 and PI19/01352, TRANSCAN EPICA AC16/00041, CIBERONC CB16/12/00489; Redes de Investigación Cooperativa (TERCEL RD16/0011/0005); Spanish Ministry of Economy, Industry and Competitivity (RTHALMY SAF2017-92632-EXP); Departamento de Salud-Gobierno de Navarra 40/2016 and Departamento de Desarrollo Económico y Empresarial (AGATA 0011-1411-2020-000010 and 0011-1411-2020-000013). The study was also supported by Cancer Research UK [C355/A26819] and Cancer Research UK, FCAECC and AIRC under the Accelerator Award Program (EDITOR), the Multiple Myeloma Research Foundation Networks of excellence 2017 Immunotherapy Program Grant Award, the International Myeloma Foundation (Brian van Novis), and Paula and Rodger Riney Foundation to FP. Instituto de Salud Carlos III (ISCIII) (PI17/01346 and PI20/00152), co-funded by the ERDF *(A way to make Europe)*; FC-AECC (AIO16163636SAEZ); Gobierno de Navarra (0011-3638-2020-000011) co-funded by the ERDF through the *Operative Program 2014-2020 of Navarra* and Gobierno de Navarra (0011-3597-2020-000005) to BS. G.T. was supported by the Associazione Italiana per la Ricerca sul Cancro (AIRC), 5×1000 Program (Project #21267). IC was supported by Juan de la Cierva grant from Ministerio de Ciencia, Innovación y Universidades, Gobierno de España, Sara Borrell award from Instituto de Salud Carlos III (ISCIII) and Marie Curie grant (H2020-MSCA-IF-837491) from European Commission. ICA was supported by AECC predoctoral fellowship.

## Data Availability

Public data was gathered from the following accessing numbers: samples GSM3674224, GSM3674225, GSM3674226, GSM3674227, GSM3674228, and GSM3674229 from GSE12842; samples GSM2915575, GSM2915576, and GSM2915577 from GSE108891. Additionally, we profiled an in-house mouse bone marrow niche sample; our raw sequencing data and expression-count will be deposited in public repositories before the publication of the manuscript. Currently, we are setting a GEO submission.

## Ethics Approval

All animal experiments were performed in accordance with national and institutional guidelines and procedures were approved by the Ethical Committee for Animal Testing of the University of Navarra. Acknowledging the principles of 3Rs (Replacement, Reduction and Refinement), all mice used in this study were from mice that were euthanized by cervical dislocation as parts of other ongoing ethically approved experiments.

The human sample collection and research conducted in this study were approved by the Research Ethics Committee of the University of Navarra. All the protocols used in this study were in strict compliance with the legal and ethical regulations.

## Author Contribution

B.S., J.T., F.P., and D.G.C. conceived and directed the research project. I.A.C. and B.S. designed the research experiments. I.A.C. with help from I.C. performed the isolation and FACS sorting of BM cells. J.Y. performed all the computational experiments, and analyzed the data with help from X.M.M., M.L. and N.P.. A.V. performed the scRNA-Seq experiments. B.S. and D.G.C. supervised the work. All authors actively participated in the discussions underlying this manuscript. I.A.C., I.C., J.Y., B.S., and D.G.C. discussed the results and wrote the final manuscript with input from J.T. and F.P.. All authors contributed to, read and approved the final manuscript.

## Competing interests

The authors declare that they have no competing interests

## Notes

### Competing Interest Statement

The authors have declared no competing interest.

### Summary of Updates

Some modifications on the text - specially in conclusions and methods are included. Additionally a link to a Github is included

## REFERENCES

Amable PR, Teixeira MVT, Carias RBV, Granjeiro JM, Borojevic R. 2014. Gene expression and protein secretion during human mesenchymal cell differentiation into adipogenic cells. BMC Cell Biology 15. doi:10.1186/s12860-014-0046-0

Aoki K, Kurashige M, Ichii M, Higaki K, Sugiyama T, Kaito T, Ando W, Sugano N, Sakai T, Shibayama H, Takaori-Kondo A, Morii E, Kanakura Y, Nagasawa T. 2021. Identification of CXCL12-abundant reticular cells in human adult bone marrow. British Journal of Haematology 193:659–668. doi:10.1111/bjh.17396

Aran D, Looney AP, Liu L, Wu E, Fong V, Hsu A, Chak S, Naikawadi RP, Wolters PJ, Abate AR, Butte AJ, Bhattacharya M. 2019. Reference-based analysis of lung single-cell sequencing reveals a transitional profibrotic macrophage. Nature Immunology 2019 20:2 20:163–172. doi:10.1038/s41590-018-0276-y

Argelaguet R, Cuomo ASE, Stegle O, Marioni JC. 2021. Computational principles and challenges in single-cell data integration. Nature Biotechnology 2021 1–14. doi:10.1038/s41587-021-00895-7

Ashhurst TM, Marsh-Wakefield F, Putri GH, Spiteri AG, Shinko D, Read MN, Smith AL, King NJC. 2021. Integration, exploration, and analysis of high-dimensional single-cell cytometry data using Spectre. Cytometry Part A. doi:10.1002/CYTO.A.24350

Baccin C, Al-Sabah J, Velten L, Helbling PM, Grünschläger F, Hernández-Malmierca P, Nombela-Arrieta C, Steinmetz LM, Trumpp A, Haas S. 2020. Combined single-cell and spatial transcriptomics reveal the molecular, cellular and spatial bone marrow niche organization. Nature Cell Biology 22:38–48. doi:10.1038/s41556-019-0439-6

Barreiro O, Yáñez-Mó M, Sala-Valdés M, Gutiérrez-López MD, Ovalle S, Higginbottom A, Monk PN, Cabañas C, Sánchez-Madrid F. 2005. Endothelial tetraspanin microdomains regulate leukocyte firm adhesion during extravasation. Blood 105:2852–2861. doi:10.1182/blood-2004-09-3606

Barrientos T, Frank D, Kuwahara K, Bezprozvannaya S, Pipes GCT, Bassel-Duby R, Richardson JA, Katus HA, Olson EN, Frey N. 2007. Two novel members of the ABLIM protein family, ABLIM-2 and -3, associate with STARS and directly bind F-actin. Journal of Biological Chemistry 282:8393–8403. doi:10.1074/jbc.M607549200

Baryawno N, Przybylski D, Kowalczyk MS, Kfoury Y, Severe N, Gustafsson K, Kokkaliaris KD, Mercier F, Tabaka M, Hofree M, Dionne D, Papazian A, Lee D, Ashenberg O, Subramanian A, Vaishnav ED, Rozenblatt-Rosen O, Regev A, Scadden DT. 2019. A Cellular Taxonomy of the Bone Marrow Stroma in Homeostasis and Leukemia. Cell 177:1915-1932.e16. doi:10.1016/j.cell.2019.04.040

Blondel VD, Guillaume J-L, Lambiotte R, Lefebvre E. 2008. Fast unfolding of communities in large networks. Journal of Statistical Mechanics: Theory and Experiment 2008:P10008. doi:10.1088/1742-5468/2008/10/P10008

Chen Q, Liu Y, Jeong HW, Stehling M, Dinh VV, Zhou B, Adams RH. 2019. Apelin+ Endothelial Niche Cells Control Hematopoiesis and Mediate Vascular Regeneration after Myeloablative Injury. Cell Stem Cell 25:768-783.e6. doi:10.1016/j.stem.2019.10.006

Chen YC, Suresh A, Underbayev C, Sun C, Singh K, Seifuddin F, Wiestner A, Pirooznia M. 2019. IKAP-Identifying K mAjor cell Population groups in single-cell RNA-sequencing analysis. GigaScience 8. doi:10.1093/gigascience/giz121

Davis GE, Senger DR. 2005. Endothelial extracellular matrix: Biosynthesis, remodeling, and functions during vascular morphogenesis and neovessel stabilization. Circulation Research. doi:10.1161/01.RES.0000191547.64391.e3

Echavarria R, Hussain SNA. 2013. Regulation of angiopoietin-1/Tie-2 receptor signaling in endothelial cells by dual-specificity phosphatases 1, 4, and 5. Journal of the American Heart Association 2. doi:10.1161/JAHA.113.000571

Escudero-Esparza A, Jiang WG, Martin TA. 2012. Claudin-5 participates in the regulation of endothelial cell motility. Molecular and Cellular Biochemistry 362:71–85. doi:10.1007/s11010-011-1129-2

Fang S, Chen S, Nurmi H, Leppänen VM, Jeltsch M, Scadden D, Silberstein L, Mikkola H, Alitalo K. 2020. VEGF-C protects the integrity of the bone marrow perivascular niche in mice. Blood 136:1871–1883. doi:10.1182/BLOOD.2020005699

Ghazanfari R, Li H, Zacharaki D, Lim HC, Scheding S. 2016. Human Non-Hematopoietic CD271pos/CD140alow/neg Bone Marrow Stroma Cells Fulfill Stringent Stem Cell Criteria in Serial Transplantations. Stem Cells and Development 25:1652–1658. doi:10.1089/scd.2016.0169

Giladi A, Paul F, Herzog Y, Lubling Y, Weiner A, Yofe I, Jaitin D, Cabezas-Wallscheid N, Dress R, Ginhoux F, Trumpp A, Tanay A, Amit I. 2018. Single-cell characterization of haematopoietic progenitors and their trajectories in homeostasis and perturbed haematopoiesis. Nature Cell Biology 2018 20:7 20:836–846. doi:10.1038/s41556-018-0121-4

Gomez-Cabrero D, Menche J, Cano I, Abugessaisa I, Huertas-Migueláñ M, Tenyi A, de Mas IM, Kiani NA, Marabita F, Falciani F, Burrowes K, Maier D, Wagner P, Selivanov V, Cascante M, Roca J, Barabási AL, Tegnér J. 2014. Systems Medicine: From molecular features and models to the clinic in COPD. Journal of Translational Medicine 12:1–11. doi:10.1186/1479-5876-12-S2-S4

Guo W, Imai S, Yang J Le, Zou S, Watanabe M, Chu YX, Mohammad Z, Xu H, Moudgil KD, Wei F, Dubner R, Ren K. 2017. In vivo immune interactions of multipotent stromal cells underlie their long-lasting pain-relieving effect. Scientific Reports 7:1–2. doi:10.1038/s41598-017-10251-y

Hafemeister C, Satija R. 2019. Normalization and variance stabilization of single-cell RNA-seq data using regularized negative binomial regression. bioRxiv 576827. doi:10.1101/576827

Hajjawi MOR, MacRae VE, Huesa C, Boyde A, Millán JL, Arnett TR, Orriss IR. 2014. Mineralisation of collagen rich soft tissues and osteocyte lacunae in Enpp1-/-mice. Bone 69:139–147. doi:10.1016/j.bone.2014.09.016

Hamidouche Z, Fromigué O, Ringe J, Häupl T, Vaudin P, Pagès JC, Srouji S, Livne E, Marie PJ. 2009. Priming integrin α5 promotes human mesenchymal stromal cell osteoblast differentiation and osteogenesis. Proceedings of the National Academy of Sciences of the United States of America 106:18587–18591. doi:10.1073/pnas.0812334106

Hao Y, Hao S, Andersen-Nissen E, Mauck WM, Zheng S, Butler A, Lee MJ, Wilk AJ, Darby C, Zager M, Hoffman P, Stoeckius M, Papalexi E, Mimitou EP, Jain J, Srivastava A, Stuart T, Fleming LM, Yeung B, Rogers AJ, McElrath JM, Blish CA, Gottardo R, Smibert P, Satija R. 2021. Integrated analysis of multimodal single-cell data. Cell 184:3573-3587.e29. doi:10.1016/J.CELL.2021.04.048

Hashemi S, Trudel E, Ganz PR, Drouin J, Couture R, Page D, Aye MT. 1991. Characterization of novel platelet and endothelial cell target antigens in a family with genetic susceptibility to autoimmunity. American Journal of Hematology 38:293–303. doi:10.1002/ajh.2830380408

Hooper AT, Butler JM, Nolan DJ, Kranz A, Iida K, Kobayashi M, Kopp HG, Shido K, Petit I, Yanger K, James D, Witte L, Zhu Z, Wu Y, Pytowski B, Rosenwaks Z, Mittal V, Sato TN, Rafii S. 2009. Engraftment and Reconstitution of Hematopoiesis Is Dependent on VEGFR2-Mediated Regeneration of Sinusoidal Endothelial Cells. Cell Stem Cell 4:263– 274. doi:10.1016/j.stem.2009.01.006

Huang W, Yang S, Shao J, Li YP. 2007. Signaling and transcriptional regulation in osteoblast commitment and differentiation. Frontiers in Bioscience 12:3068–3092. doi:10.2741/2296

Itkin T, Gur-Cohen S, Spencer JA, Schajnovitz A, Ramasamy SK, Kusumbe AP, Ledergor G, Jung Y, Milo I, Poulos MG, Kalinkovich A, Ludin A, Kollet O, Shakhar G, Butler JM, Rafii S, Adams RH, Scadden DT, Lin CP, Lapidot T. 2016. Distinct bone marrow blood vessels differentially regulate haematopoiesis. Nature 532:323–328. doi:10.1038/nature17624

Jia J, Ye T, Cui P, Hua Q, Zeng H, Zhao D. 2016. AP-1 transcription factor mediates VEGF-induced endothelial cell migration and proliferation. Microvascular Research 105:103–108. doi:10.1016/j.mvr.2016.02.004

Kahai S, Vary CPH, Gao Y, Seth A. 2004. Collagen, type V, α1 (COL5A1) is regulated by TGF-β in osteoblasts. Matrix Biology 23:445–455. doi:10.1016/j.matbio.2004.09.004

Kalucka J, de Rooij LPMH, Goveia J, Rohlenova K, Dumas SJ, Meta E, Conchinha N V., Taverna F, Teuwen LA, Veys K, García-Caballero M, Khan S, Geldhof V, Sokol L, Chen R, Treps L, Borri M, de Zeeuw P, Dubois C, Karakach TK, Falkenberg KD, Parys M, Yin X, Vinckier S, Du Y, Fenton RA, Schoonjans L, Dewerchin M, Eelen G, Thienpont B, Lin L, Bolund L, Li X, Luo Y, Carmeliet P. 2020. Single-Cell Transcriptome Atlas of Murine Endothelial Cells. Cell 180:764-779.e20. doi:10.1016/j.cell.2020.01.015

Kanazawa S, Okada H, Hojo H, Ohba S, Iwata J, Komura M, Hikita A, Hoshi K. 2021. Mesenchymal stromal cells in the bone marrow niche consist of multi-populations with distinct transcriptional and epigenetic properties. Scientific Reports 11. doi:10.1038/s41598-021-94186-5

Karamitros D, Stoilova B, Aboukhalil Z, Hamey F, Reinisch A, Samitsch M, Quek L, Otto G, Repapi E, Doondeea J, Usukhbayar B, Calvo J, Taylor S, Goardon N, Six E, Pflumio F, Porcher C, Majeti R, Göttgens B, Vyas P. 2017. Single-cell analysis reveals the continuum of human lympho-myeloid progenitor cells. Nature Immunology 2017 19:1 19:85–97. doi:10.1038/s41590-017-0001-2

Kenswil KJG, Jaramillo AC, Ping Z, Chen S, Hoogenboezem RM, Mylona MA, Adisty MN, Bindels EMJ, Bos PK, Stoop H, Lam KH, van Eerden B, Cupedo T, Raaijmakers MHGP. 2018. Characterization of Endothelial Cells Associated with Hematopoietic Niche Formation in Humans Identifies IL-33 As an Anabolic Factor. Cell Reports 22:666–678. doi:10.1016/j.celrep.2017.12.070

Kim SH, In Choi H, Choi MR, An GY, Binas B, Jung KH, Chai YG. 2020. Epigenetic regulation of IFITM1 expression in lipopolysaccharide-stimulated human mesenchymal stromal cells. Stem Cell Research and Therapy 11:1–12. doi:10.1186/s13287-019-1531-3

Klóska D, Kopacz A, Piechota-Polańczyk A, Neumayer C, Huk I, Dulak J, Józkowicz A, Grochot-Przęczek A. 2019. Biliverdin reductase deficiency triggers an endothelial-to-mesenchymal transition in human endothelial cells. Archives of Biochemistry and Biophysics 678:108182. doi:10.1016/j.abb.2019.108182

Laurenti E, Göttgens B. 2018. From haematopoietic stem cells to complex differentiation landscapes. Nature. doi:10.1038/nature25022

Lee JS, Yu Q, Shin JT, Sebzda E, Bertozzi C, Chen M, Mericko P, Stadtfeld M, Zhou D, Cheng L, Graf T, MacRae CA, Lepore JJ, Lo CW, Kahn ML. 2006. Klf2 Is an Essential Regulator of Vascular Hemodynamic Forces In Vivo. Developmental Cell 11:845–857. doi:10.1016/j.devcel.2006.09.006

Leimkühler NB, Gleitz HFE, Ronghui L, Snoeren IAM, Fuchs SNR, Nagai JS, Banjanin B, Lam KH, Vogl T, Kuppe C, Stalmann USA, Büsche G, Kreipe H, Gütgemann I, Krebs P, Banz Y, Boor P, Tai EWY, Brümmendorf TH, Koschmieder S, Crysandt M, Bindels E, Kramann R, Costa IG, Schneider RK. 2021. Heterogeneous bone-marrow stromal progenitors drive myelofibrosis via a druggable alarmin axis. Cell Stem Cell 28:637-652.e8. doi:10.1016/j.stem.2020.11.004

Linnemann AK, O’Geen H, Keles S, Farnham PJ, Bresnick EH. 2011. Genetic framework for GATA factor function in vascular biology. Proceedings of the National Academy of Sciences of the United States of America 108:13641–13646. doi:10.1073/pnas.1108440108

Mackenzie NCW, Zhu D, Milne EM, van’t Hof R, Martin A, Quarles DL, Millán JL, Farquharson C, MacRae VE. 2012. Altered bone development and an increase in FGF-23 expression in Enpp1 -/-mice. PLoS ONE 7:32177. doi:10.1371/journal.pone.0032177

Macosko EZ, Basu A, Satija R, Nemesh J, Shekhar K, Goldman M, Tirosh I, Bialas AR, Kamitaki N, Martersteck EM, Trombetta JJ, Weitz DA, Sanes JR, Shalek AK, Regev A, McCarroll SA. 2015. Highly parallel genome-wide expression profiling of individual cells using nanoliter droplets. Cell 161:1202–1214. doi:10.1016/j.cell.2015.05.002

March S, Hui EE, Underhill GH, Khetani S, Bhatia SN. 2009. Microenvironmental regulation of the sinusoidal endothelial cell phenotype in vitro. Hepatology 50:920–928. doi:10.1002/hep.23085

Martinez-de-Morentin X, Khan SA, Lehmann R, Tegner J, Gomez-Cabrero D. 2021. Machine Translation between paired Single Cell Multi Omics Data. bioRxiv 2021.01.27.428400. doi:10.1101/2021.01.27.428400

Matsushita Y, Nagata M, Kozloff KM, Welch JD, Mizuhashi K, Tokavanich N, Hallett SA, Link DC, Nagasawa T, Ono W, Ono N. 2020. A Wnt-mediated transformation of the bone marrow stromal cell identity orchestrates skeletal regeneration. Nature Communications 11. doi:10.1038/s41467-019-14029-w

Matthews E, Lanham S, White K, Kyriazi ME, Alexaki K, El-Sagheer AH, Brown T, Kanaras AG, West J, MacArthur BD, Stumpf PS, Oreffo ROC. 2020. Single cell RNA sequence analysis of human bone marrow samples reveals new targets for isolation of skeletal stem cells using DNA-coated gold nanoparticles. bioRxiv. doi:10.1101/2020.06.17.156836

Naito H, Iba T, Takakura N. 2021. Mechanisms of new blood-vessel formation and proliferative heterogeneity of endothelial cells. International Immunology 32:295–305. doi:10.1093/intimm/dxaa008

Nakamura S, Koyama T, Izawa N, Nomura S, Fujita T, Omata Y, Minami T, Matsumoto M, Nakamura M, Fujita-Jimbo E, Momoi T, Miyamoto T, Aburatani H, Tanaka S. 2017. Negative feedback loop of bone resorption by NFATc1-dependent induction of Cadm1. PLoS ONE 12:1–14. doi:10.1371/journal.pone.0175632

Nestorowa S, Hamey FK, Pijuan Sala B, Diamanti E, Shepherd M, Laurenti E, Wilson NK, Kent DG, Göttgens B. 2016. A single-cell resolution map of mouse hematopoietic stem and progenitor cell differentiation. Blood 128:e20–e31. doi:10.1182/BLOOD-2016-05-716480

Nilius B, Droogmans G. 2001. Ion channels and their functional role in vascular endothelium. Physiological Reviews. doi:10.1152/physrev.2001.81.4.1415

Okamoto T, Suzuki K. 2017. The role of gap junction-mediated endothelial cell-cell interaction in the crosstalk between inflammation and blood coagulation. International Journal of Molecular Sciences 18. doi:10.3390/ijms18112254

Price PA. 1985. Vitamin K-Dependent Formation of Bone Gla Protein (Osteocalcin) and Its Function. Vitamins and Hormones 42:65–108. doi:10.1016/S0083-6729(08)60061-8

Quillon A, Fromy B, Debret R. 2015. Endothelium microenvironment sensing leading to nitric oxide mediated vasodilation: A review of nervous and biomechanical signals. Nitric Oxide - Biology and Chemistry. doi:10.1016/j.niox.2015.01.006

Quirici N, Soligo D, Bossolasco P, Servida F, Lumini C, Deliliers GL. 2002. Isolation of bone marrow mesenchymal stem cells by anti-nerve growth factor receptor antibodies. Experimental Hematology 30:783–791. doi:10.1016/S0301-472X(02)00812-3

Raaijmakers MHGP, Mukherjee S, Guo S, Zhang S, Kobayashi T, Schoonmaker JA, Ebert BL, Al-Shahrour F, Hasserjian RP, Scadden EO, Aung Z, Matza M, Merkenschlager M, Lin C, Rommens JM, Scadden DT. 2010. Bone progenitor dysfunction induces myelodysplasia and secondary leukaemia. Nature 464:852–857. doi:10.1038/nature08851

Rafii S, Butler JM, Ding B Sen. 2016. Angiocrine functions of organ-specific endothelial cells. Nature 529:316–325. doi:10.1038/nature17040

Robert AW, Marcon BH, Dallagiovanna B, Shigunov P. 2020. Adipogenesis, Osteogenesis, and Chondrogenesis of Human Mesenchymal Stem/Stromal Cells: A Comparative Transcriptome Approach. Frontiers in Cell and Developmental Biology. doi:10.3389/fcell.2020.00561

Rodriguez-Fraticelli AE, Wolock SL, Weinreb CS, Panero R, Patel SH, Jankovic M, Sun J, Calogero RA, Klein AM, Camargo FD. 2018. Clonal analysis of lineage fate in native haematopoiesis. Nature 2018 553:7687 553:212–216. doi:10.1038/nature25168

Severe N, Karabacak NM, Gustafsson K, Baryawno N, Courties G, Kfoury Y, Kokkaliaris KD, Rhee C, Lee D, Scadden EW, Garcia-Robledo JE, Brouse T, Nahrendorf M, Toner M, Scadden DT. 2019. Stress-Induced Changes in Bone Marrow Stromal Cell Populations Revealed through Single-Cell Protein Expression Mapping. Cell Stem Cell 25:570-583.e7. doi:10.1016/j.stem.2019.06.003

Shen B, Tasdogan A, Ubellacker JM, Zhang J, Nosyreva ED, D. L, Murphy MM, Hu S, Yi Y, Kara N, Liu X, Guela S, Jia Y, Ramesh V, Embree C, Mitchell EC, Zhao YC, Ju LA, Hu Z, Crane GM, Zhao Z, Syeda R, Morrison SJ. 2021. A mechanosensitive peri-arteriolar niche for osteogenesis and lymphopoiesis. Nature 591:438–444. doi:10.1038/s41586-021-03298-5

Spohn G, Kleinridders A, Wunderlich FT, Watzka M, Zaucke F, Blumbach K, Geisen C, Seifried E, Müller C, Paulsson M, Brüning JC, Oldenburg J. 2009. VKORC1 deficiency in mice causes early postnatal lethality due to severe bleeding. Thrombosis and Haemostasis 101:1044–1050. doi:10.1160/TH09-03-0204

Stuart T, Butler A, Hoffman P, Hafemeister C, Papalexi E, Mauck WM, Hao Y, Stoeckius M, Smibert P, Satija R. 2019. Comprehensive Integration of Single-Cell Data. Cell 177:1888-1902.e21. doi:10.1016/j.cell.2019.05.031

Stumpf PS, D. X, Imanishi H, Kunisaki Y, Semba Y, Noble T, Smith RCG, Rose-Zerili M, West JJ, Oreffo ROC, Farrahi K, Niranjan M, Akashi K, Arai F, MacArthur BD. 2020. Transfer learning efficiently maps bone marrow cell types from mouse to human using single-cell RNA sequencing. Communications Biology 3. doi:10.1038/s42003-020-01463-6

Sureshbabu A, Okajima H, Yamanaka D, Tonner E, Shastri S, Maycock J, Szymanowska M, Shand J, Takahashi SI, Beattie J, Allan G, Flint D. 2012. IGFBP5 induces cell adhesion, increases cell survival and inhibits cell migration in MCF-7 human breast cancer cells. Journal of Cell Science 125:1693–1705. doi:10.1242/jcs.092882

Tamura K, Hashimoto K, Suzuki K, Yoshie M, Kutsukake M, Sakurai T. 2009. Insulin-like growth factor binding protein-7 (IGFBP7) blocks vascular endothelial cell growth factor (VEGF)-induced angiogenesis in human vascular endothelial cells. European Journal of Pharmacology 610:61–67. doi:10.1016/j.ejphar.2009.01.045

Tasic B, Menon V, Nguyen TN, Kim TK, Jarsky T, Yao Z, Levi B, Gray LT, Sorensen SA, Dolbeare T, Bertagnolli D, Goldy J, Shapovalova N, Parry S, Lee C, Smith K, Bernard A, Madisen L, Sunkin SM, Hawrylycz M, Koch C, Zeng H. 2016a. Adult mouse cortical cell taxonomy revealed by single cell transcriptomics. Nature Neuroscience 19:335–346. doi:10.1038/nn.4216

Tasic B, Menon V, Nguyen TN, Kim TK, Jarsky T, Yao Z, Levi B, Gray LT, Sorensen SA, Dolbeare T, Bertagnolli D, Goldy J, Shapovalova N, Parry S, Lee C, Smith K, Bernard A, Madisen L, Sunkin SM, Hawrylycz M, Koch C, Zeng H. 2016b. Adult mouse cortical cell taxonomy revealed by single cell transcriptomics. Nature Neuroscience 19:335–346. doi:10.1038/nn.4216

Tegnér JN, Compte A, Auffray C, An G, Cedersund G, Clermont G, Gutkin B, Oltvai ZN, Stephan KE, Thomas R, Villoslada P. 2009. Computational disease modeling – fact or fiction? BMC Systems Biology 2009 3:1 3:1–3. doi:10.1186/1752-0509-3-56

Testa U, Labbaye C, Castelli G, Pelosi E. 2016. Oxidative stress and hypoxia in normal and leukemic stem cells. Experimental Hematology. doi:10.1016/j.exphem.2016.04.012

Tikhonova AN, Dolgalev I, Hu H, Sivaraj KK, Hoxha E, Cuesta-Domínguez Á, Pinho S, Akhmetzyanova I, Gao J, Witkowski M, Guillamot M, Gutkin MC, Zhang Y, Marier C, Diefenbach C, Kousteni S, Heguy A, Zhong H, Fooksman DR, Butler JM, Economides A, Frenette PS, Adams RH, Satija R, Tsirigos A, Aifantis I. 2019. The bone marrow microenvironment at single-cell resolution. Nature 569:222–228. doi:10.1038/s41586-019-1104-8

Traag VA, Waltman L, Eck NJ van. 2019. From Louvain to Leiden: guaranteeing well-connected communities. Scientific Reports 2019 9:1 9:1–12. doi:10.1038/s41598-019-41695-z

Velten L, Haas SF, Raffel S, Blaszkiewicz S, Islam S, Hennig BP, Hirche C, Lutz C, Buss EC, Nowak D, Boch T, Hofmann W-K, Ho AD, Huber W, Trumpp A, Essers MAG, Steinmetz LM. 2017. Human haematopoietic stem cell lineage commitment is a continuous process. Nature Cell Biology 2017 19:4 19:271–281. doi:10.1038/ncb3493

Wang Z, Li X, Yang J, Gong Y, Zhang H, Qiu X, Liu Y, Zhou C, Chen Y, Greenbaum J, Cheng L, Hu Y, Xie J, Yang X, Li Y, Schiller M, Tan L, Tang S-Y, Shen H, Xiao H-M, Deng H-W. 2020. Single-cell RNA sequencing deconvolutes the in vivo heterogeneity of human bone marrow-derived mesenchymal stem cells. bioRxiv 2020.04.06.027904. doi:10.1101/2020.04.06.027904

Weinreb C, Rodriguez-Fraticelli A, Camargo FD, Klein AM. 2020. Lineage tracing on transcriptional landscapes links state to fate during differentiation. Science 367. doi:10.1126/SCIENCE.AAW3381

Wolkenhauer O, Auffray C, Jaster R, Steinhoff G, Dammann O. 2013. The road from systems biology to systems medicine. Pediatric Research. doi:10.1038/pr.2013.4

Wolock SL, Krishnan I, Tenen DE, Matkins V, Camacho V, Patel S, Agarwal P, Bhatia R, Tenen DG, Klein AM, Welner RS. 2019. Mapping Distinct Bone Marrow Niche Populations and Their Differentiation Paths. Cell Reports 28:302-311.e5. doi:10.1016/j.celrep.2019.06.031

Wu KE, Yost KE, Chang HY, Zou J. n.d. BABEL enables cross-modality translation between multiomic profiles at single-cell resolution. doi:10.1073/pnas.2023070118/-/DCSupplemental

Xie J, Lou Q, Zeng Y, Liang Y, Xie S, Xu Q, Yuan L, Wang J, Jiang L, Mou L, Lin D, Zhao M. 2021. Single-Cell Atlas Reveals Fatty Acid Metabolites Regulate the Functional Heterogeneity of Mesenchymal Stem Cells. Frontiers in Cell and Developmental Biology 9:1–15. doi:10.3389/fcell.2021.653308

Yau JW, Teoh H, Verma S. 2015. Endothelial cell control of thrombosis. BMC Cardiovascular Disorders 15:130. doi:10.1186/s12872-015-0124-z

Yu G, Wang LG, Han Y, He QY. 2012. ClusterProfiler: An R package for comparing biological themes among gene clusters. OMICS A Journal of Integrative Biology 16:284–287. doi:10.1089/omi.2011.0118

Yue R, Shen B, Morrison SJ. 2016. Clec11a/osteolectin is an osteogenic growth factor that promotes the maintenance of the adult skeleton. eLife 5:27. doi:10.7554/eLife.18782

Zambetti NA, Ping Z, Chen S, Kenswil KJG, Mylona MA, Sanders MA, Hoogenboezem RM, Bindels EMJ, Adisty MN, Van Strien PMH, van der Leije CS, Westers TM, Cremers EMP, Milanese C, Mastroberardino PG, van Leeuwen JPTM, van der Eerden BCJ, Touw IP, Kuijpers TW, Kanaar R, van de Loosdrecht AA, Vogl T, Raaijmakers MHGP. 2016. Mesenchymal Inflammation Drives Genotoxic Stress in Hematopoietic Stem Cells and Predicts Disease Evolution in Human Pre-leukemia. Cell Stem Cell 19:613–627. doi:10.1016/j.stem.2016.08.021

Zhao E, Xu H, Wang L, Kryczek I, Wu K, Hu Y, Wang G, Zou W. 2012. Bone marrow and the control of immunity. Cellular and Molecular Immunology. doi:10.1038/cmi.2011.47

Zhong L, Yao L, Tower RJ, Wei Y, Miao Z, Park J, Shrestha R, Wang L, Yu W, Holdreith N, Huang X, Zhang Y, Tong W, Gong Y, Ahn J, Susztak K, Dyment N, Li M, Long F, Chen C, Seale P, Qin L. 2020. Single cell transcriptomics identifies a unique adipose lineage cell population that regulates bone marrow environment. eLife 9:1–28. doi:10.7554/eLife.54695

