## Supplementary material for "Deep deconvolution of the hematopoietic stem cell regulatory microenvironment reveals a high degree of specialization and conservation between mouse and human": Supp. Information.

#### Supplementary Figures

**Figure S1. Cell sorting strategy for in-house dataset.** Cell sorting strategy for isolation of mouse BM microenvironment cells (Lived niche cells: Annexin V$^-$ 7-AAD$^-$ Lin$^-$ CD45$^-$ Ter119$^-$ VDO$^+$).

**Figure S2. Additional information on the clustering analysis. (a)** Description of the bootstrapping strategy for evaluation cluster robustness (see also Methods). **(b)** Representation of the number of cells for each cluster and contribution of each dataset to the final endothelial clusters. **(c-e)** Illustration of the sub-clustering, recall per cell and summary of recall per cells for every cluster respectively for the analysis of cluster A1. **(f-h)** Demonstration of the same analysis for cluster A2.

**Figure S3. High-resolution clustering strategy of mesenchymal cells. (a)** Clustering strategy: Mesenchymal. An upper limit to cluster is set for the clustering (left panel) using Louvain high-resolution clustering. Then, an iterative divide-and-conquer strategy identifies the optimal level of clusters for a given set of cells, and, within each cluster, it evaluates possible sub-clustering. Level 1 (second panel from the left), Level 2 (third panel) and Level 3 (fourth panel) shows the clustering after every iteration. **(b)** Representation of the number of cells for each cluster and contribution of each data set to final mesenchymal clusters. **(c-e)** Robustness of the analysis for sub-clustering C2 (from Level 2 to Level 3). Specifically: **(c)** Illustration of the identified clusters**; (d)** Depiction of every cell, *how many times it is assigned correctly to a cluster using a random-forest + bootstrapping strategy* (see Methods); **(e)** Summary of the results (d) per cluster. **(f-h)** Demonstration of the same analysis for cluster C4.

**Figure S4. Added value: identification of clusters**. Pairwise Jaccard Index analysis between Baryawno dataset only and the identified clusters observed with the integrated dataset (rows).

**Figure S5. Analysis of human samples. (a)** Cell sorting strategy for isolation of human bone marrow endothelial (TO-PRO-3$^-$, Lin$^-$, CD45$^{-,}$ CD235$^-$, CD9$^+$, CD31$^+$) and mesenchymal-osteolineage (TO-PRO-3$^-$, Lin$^-$, CD45$^-$, CD235$^-$, CD31$^-$, CD271$^+$, CD146$^{+/-}$) cells. **(b)** Left panel: UMAP representation of cells quantified after filtering showing the sample of origin; Middle Panel: number of cells per cluster and sample of origin; Right panel: proportion of cells per cluster and sample of origin. **(c)** Left panel: UMAP visualization of cells quantified after filtering and showing the cell cycle stage; Middle Panel: number of cells per cluster and cell cycle stage; Right panel: proportion of cells per cluster and cell cycle stage.

**Figure S6. Conservation analysis of the EC and MSC population in the human BM microenvironment.** Annotation of the human cells using the identified endothelial (a) and mesenchymal (c) mouse markers through SingleR analysis. Violin plots of the associated scores for endothelial (b) and mesenchymal (d) subclusters.

**Figure S7. Assignment of cells from non-robust clusters.** Example of how cells can be assigned to other clusters once they have been identified as non-robust.

### Supplementary Figures

Table S1. Marker genes per cluster in Mouse endothelial cells (EC).
Table S2. Marker genes between sinusoids and arteries in Mouse endothelial cells (EC).
Table S3. Gene-Set Enrichment per cluster in Mouse endothelial cells (EC).
Table S4. Marker genes per cluster in Mouse Mesenchymal cells (MSC).
Table S5. Marker genes between mesenchymal and osteolineage (OLN-primed) in Mouse Mesenchymal cells (MSC).
Table S6. Gene-Set Enrichment per cluster in Mouse Mesenchymal cells (MSC).
Table S7. Human Cluster Marker after IG Filtering.
Table S8. Gene Set Enrichment in Human integrated data.
Table S9. Genes from mouse endothelial (EC) clusters in Human data.
Table S10. Genes from mouse mesenchymal (MSC) clusters in Human data.
