## Supplementary material for "Deep deconvolution of the hematopoietic stem cell regulatory microenvironment reveals a high degree of specialization and conservation between mouse and human": Supp. Figures

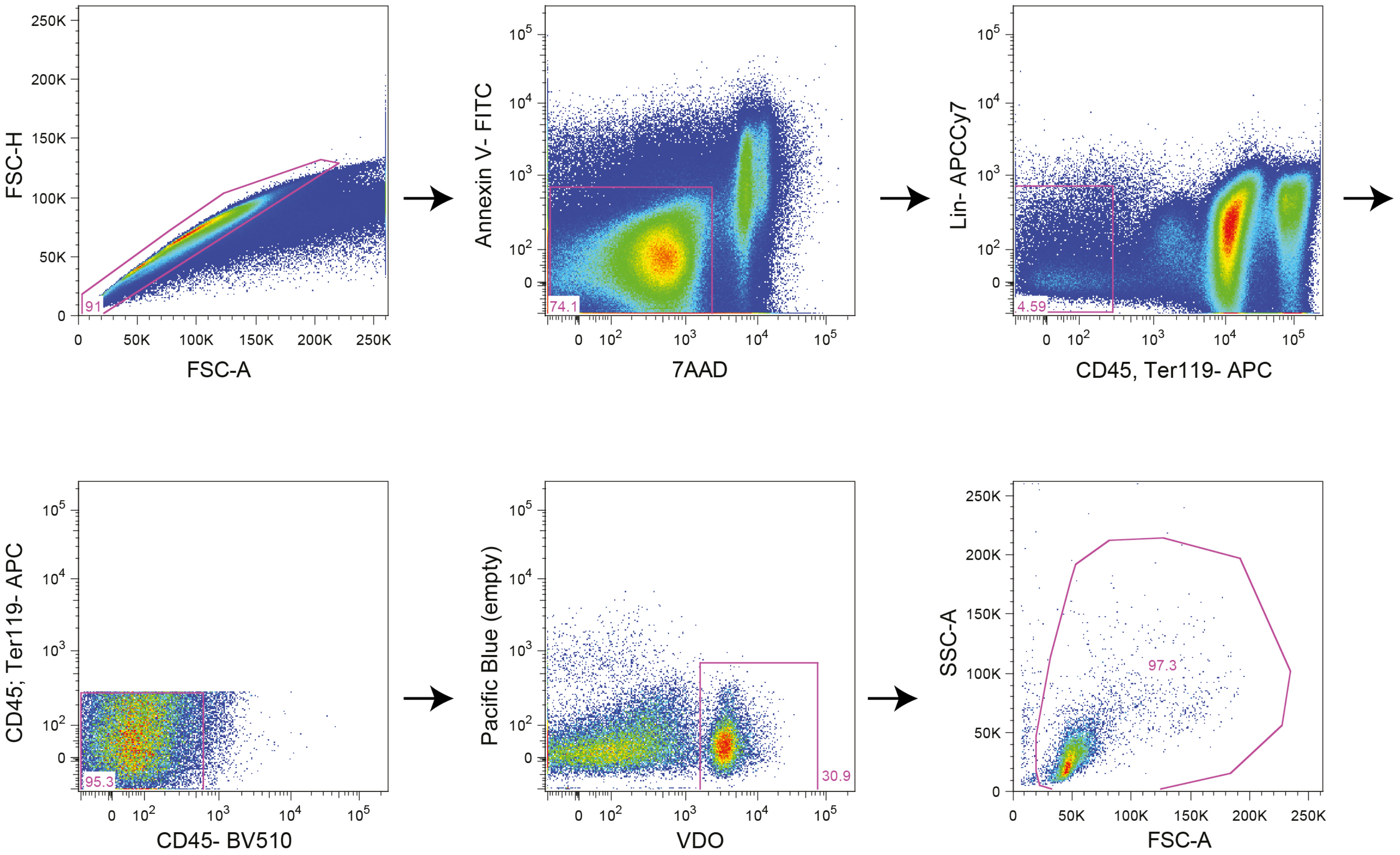


Ye et al, Figure S1

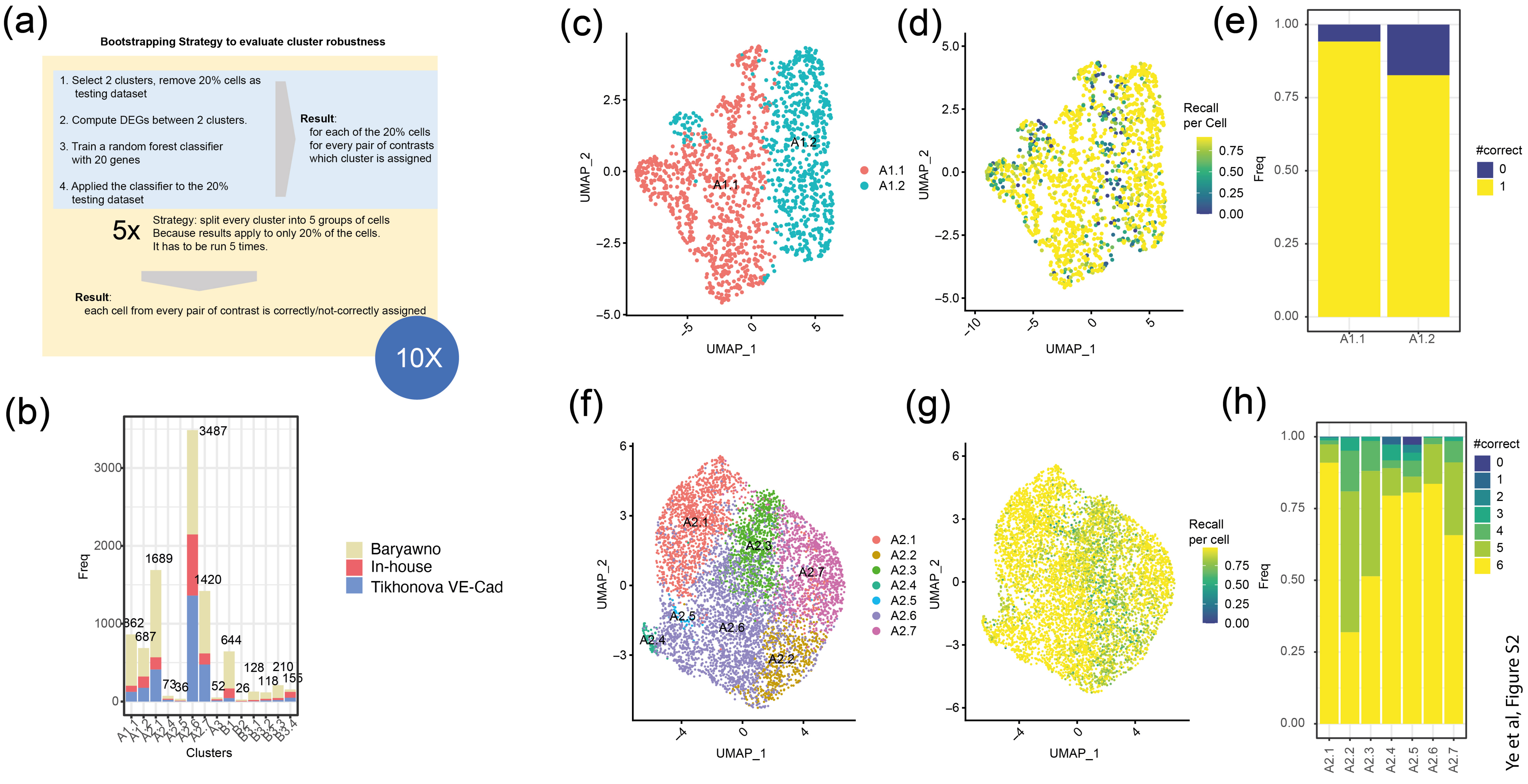


Ye et al, Figure S2

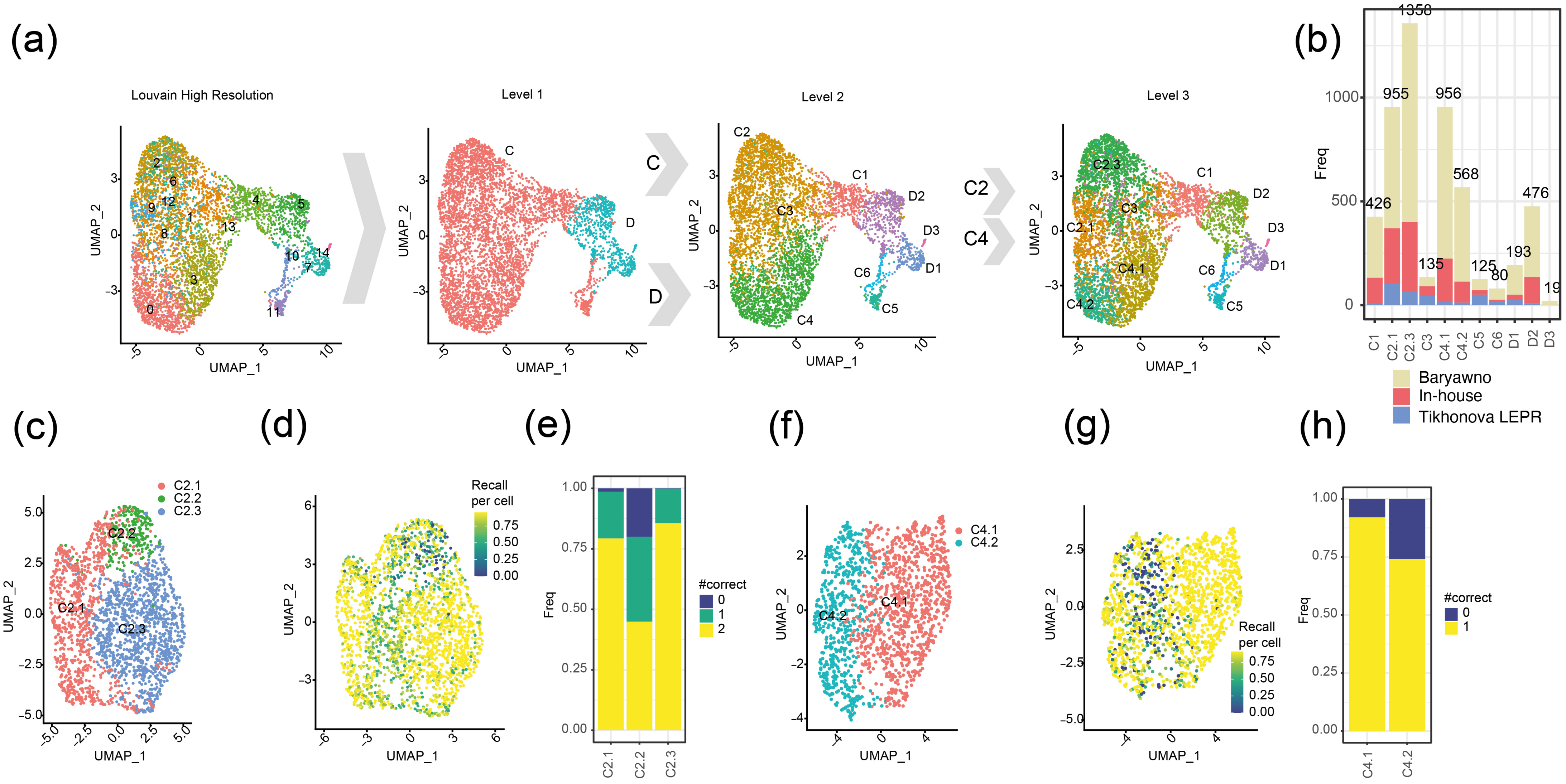


Ye et al, Figure S3

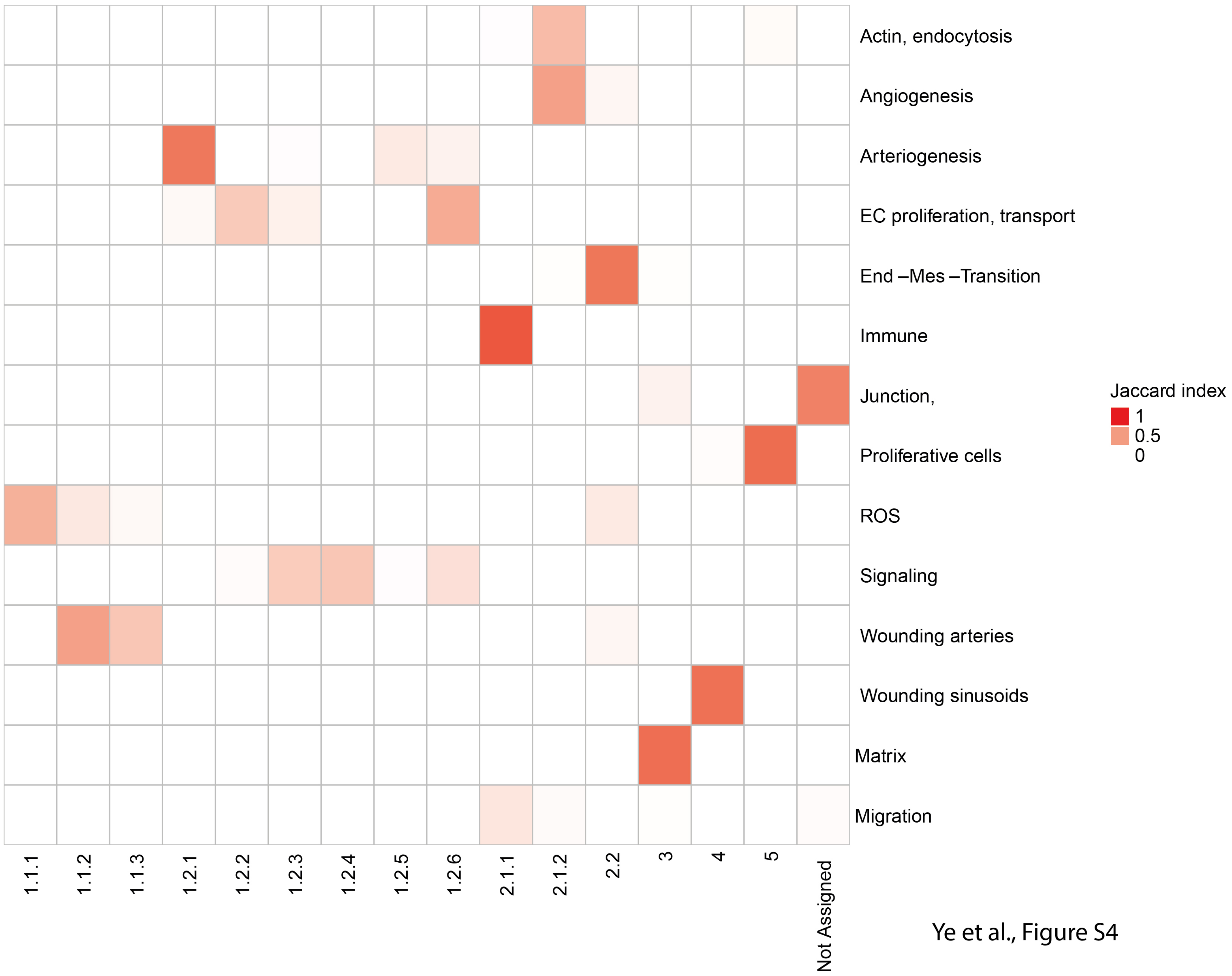


Ye et al., Figure S4

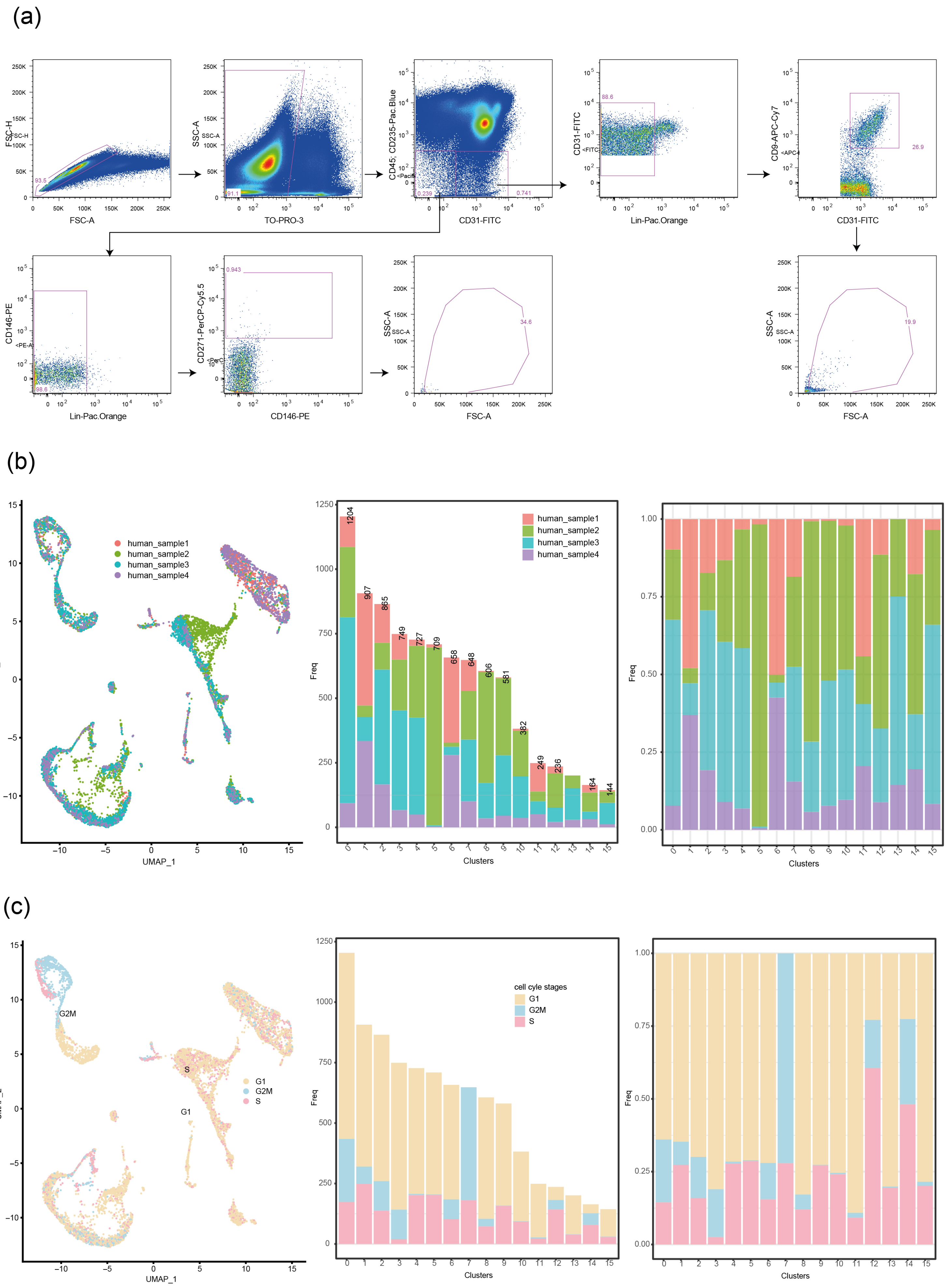


Ye et al., Figure S5

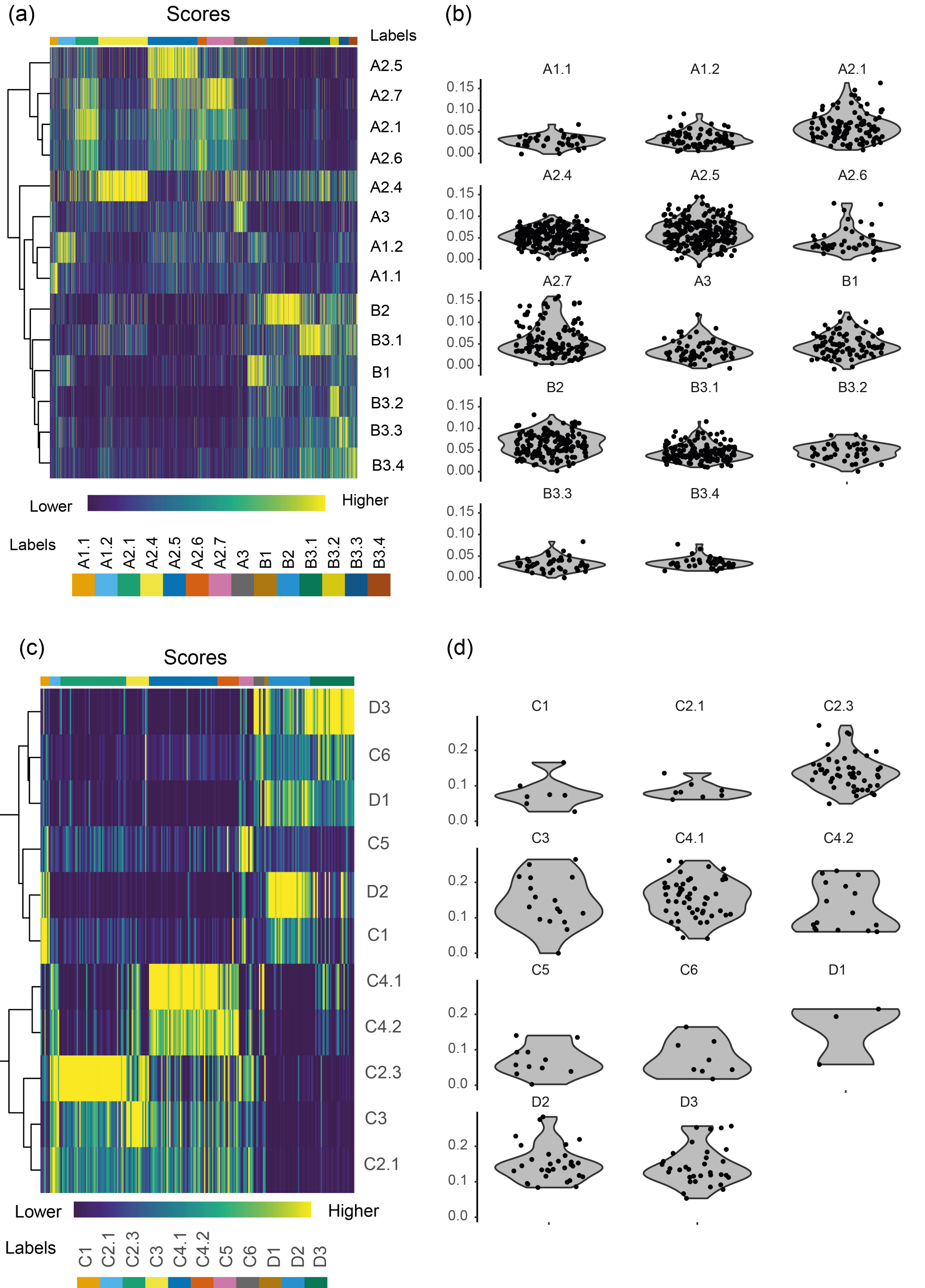


Ye et al, Figure S6

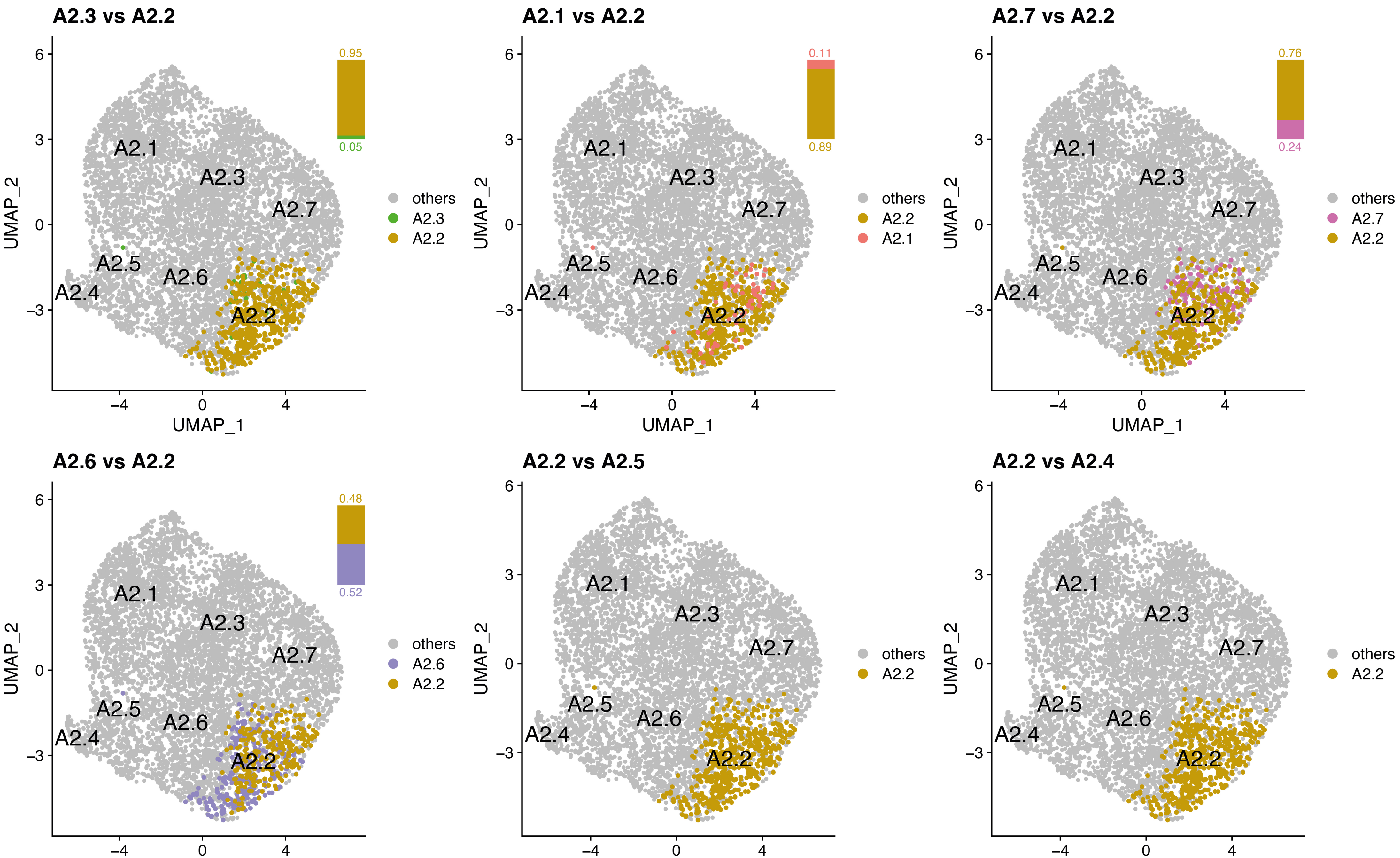


Ye et al, Figure S7
